# Hotspots of pest-induced US urban tree death, 2020-2050

**DOI:** 10.1101/2021.04.24.441210

**Authors:** Emma J. Hudgins, Frank H. Koch, Mark J. Ambrose, Brian Leung

## Abstract

1. Urban trees are important nature-based solutions for future wellbeing and livability but are at high risk of mortality from insect pests. In the United States (US), 82% of the population live in urban settings and this number is growing, making urban tree mortality a matter of concern for most of its population. Until now, the magnitudes and spatial distributions of risks were unknown.
2. Here, we combine new models of street tree populations in ∼30,000 US communities, species-specific spread predictions for 57 invasive insect species, and estimates of tree death due to insect exposure for 48 host tree genera.
3. We estimate that 1.4 million street trees will be killed by invasive insects from 2020 through 2050, costing an annualized average of US$ 30M. However, these estimates hide substantial variation: 23% of urban centers will experience 95% of all insect-induced mortality. Further, 90% of all mortality will be due to emerald ash borer (*Agrilus planipennis*, EAB), which is expected to kill virtually all ash trees (*Fraxinus* spp.) in >6000 communities.
4. We define an EAB high-impact zone spanning 902,500km^2^, largely within the southern and central US, within which we predict the death of 98.8% of all ash trees. “Mortality hotspot cities” include Milwaukee, WI; Chicago, IL; and New York, NY.
5. We identify Asian wood borers of maple and oak trees as the highest risk future invaders, where a new establishment could cost US$ 4.9B over 30 years.
6. *Policy implications:* To plan effective mitigation, managers must know which tree species in which communities will be at the greatest risk, as well as the highest-risk insects. We provide the first country-wide, spatial forecast of urban tree mortality due to invasive insect pests. This framework identifies dominant pest insects and spatial impact hotspots, which can provide the basis for spatial prioritization of spread control efforts such as quarantines and biological control release sites. Our results highlight the need for EAB early-detection efforts as far from current infestations as Seattle, WA. Further, these findings produce a list of biotic and spatiotemporal risk factors for future high-impact US urban forest insect pests.

## Introduction

Urban tree impacts comprise the dominant share of economic damages caused by invasive alien forest insects (IAFIs) in the United States (US) (Aukema et al. 2011; Holmes et al. 2009; Kovacs et al. 2010; Lovett et al. 2016). Urban trees are known to facilitate the invasion of IAFIs (Branco et al. 2019), and urban forests located close to points of IAFI entry are often the first establishment locations of new IAFIs (Rassati et al. 2015). Further, urban tree populations include highly susceptible species such as ash (*Fraxinus* spp.) that are being decimated by emerald ash borer (EAB, *Agrilus planipennis*) (Kovacs et al. 2010). To eliminate the potential for injury or property damage due to dead trees, infested urban trees must be treated or removed (Fahrner et al. 2017). Moreover, the importance of urban trees is only expected to grow. While the percentage of people living in cities is already very high in the US (82% in 2018), it has not yet peaked (World Bank, http://data.worldbank.org, UN DESA, https://population.un.org/wup/). At the same time, there has been a push for urban ‘greening’ (i.e., increasing urban tree canopy). Urban trees provide many important ecosystem services, including lowering cooling costs (Norton et al. 2015), buffering against flooding, improving air quality, carbon sequestration, improving citizens’ mental and physical health outcomes, and creating important wildlife habitat (Van den Berg et al. 2010; Roy et al. 2012). The high tree mortality risk posed by IAFIs can greatly diminish these myriad benefits.

While IAFI life histories differ, they are known to be transported long distances by humans (Hulme 2009), potentially with similar drivers across entire secondary dispersal pathways following establishment within a country (Hudgins et al. 2017, 2020). Thus, the creation of a pathway-level damage estimate can provide insight into the benefit of limiting future spread via these pathways (e.g., through quarantines or roadside checkpoints to limit firewood movement). Past estimates of IAFI damage have been important in providing support for phytosanitary measures such as ISPM15 (IPPC 2002), a wood packing material treatment protocol, whose adoption is growing worldwide (Leung et al. 2014). A previous pathway-level estimate for the cumulative cost of all US IAFIs was performed a decade ago but had substantial data limitations (Aukema et al. 2011), including reliance on detailed dispersal projections for only three species, tree distributions based on only a handful of US cities, and spatially implicit impact forecasts. Since then, contemporary advances in modelling and increased data availability allow direct estimates of spread for every IAFI species as well as host prevalence and IAFI-induced mortality for every tree species in every community across the United States. This enables not only the estimation of country-wide IAFI damages, but also IAFI and host-specific damages and their spatial distribution. Additionally, we can examine the impact of tree mortality dynamics on cost dynamics and derive better risk assessments of not-yet established pests, based on their functional traits and host distributions.

In this paper, we synthesized four subcomponents of IAFI invasions: 1) a model of spread for 57 IAFI species, 2) a model for the distribution of all urban street tree host genera across all US communities, 3) a model of host mortality in response to IAFI-specific infestation for all urban host tree species, and 4) the cost of removing and replacing dead trees, to provide the best current estimate of the damage to street trees, including explicit estimates for all known IAFIs across all major insect guilds. With this model, we aimed to determine the US communities, host tree species, and pest species associated with the greatest forest pest impacts in the next 30 years. From these results, we wished to distill a set of high-risk factors for future IAFIs not currently established in the US.

## Materials and Methods

We synthesized four subcomponent models of IAFI invasions (see conceptual diagram, Fig. S1).

### IAFI dispersal forecasts

We modelled spread using the Semi-Generalized Dispersal Kernel (SDK, Hudgins et al. 2020). This is a spatially explicit, negative exponential dispersal kernel model fit from a simulation procedure that can account for additional spatial predictors in source and recipient sites by treating them as factors that increase or decrease dispersal into or out of these sites. At each timestep, pests disperse according to the following equation,

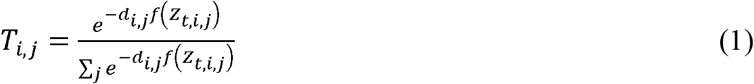

where *i* and *j* correspond to source and recipient simulation grid cells, *d* corresponds to the Euclidean distance between sites *i* and *j* and *f*(*Z*_*t,i,j*_) corresponds to a logistically-scaled set of spatial predictors in source and destination sites at a particular timestep *t*.

Importantly, only a single snapshot of pest species distributions is required to fit the model. The SDK builds from the Generalized Dispersal Kernel (GDK, Hudgins et al. 2017) as a starting point, using human population density, forested land area and tree density in source and destination sites (i.e., simulation model cells donating and receiving propagules to or from other cells) as moderators of spread. The SDK combines up to three species-specific corrections for each species to maximize predictive ability: 1) a species-specific intercept term; 2) information on an IAFI’s likely initial invasion location derived from expert opinion; and 3) niche-related limitations when evidenced in the literature, including a fitted minimum January temperature threshold for hemlock woolly adelgid (*Adelges tsugae*). The SDK was applied to 57 IAFIs believed to cause some damage from Aukema et al. (2011, Table S1) and projected from 2020 to 2050 (Fig. S2). This list excludes species with less than five years of history in the US, any species known to already have reasonable levels of spread control (i.e., *Anoplophora glabripennis*), species without known economic impacts, and species lacking known host distributions in natural forests (Liebhold et al. 2013),

### Street tree models

Primarily, our analysis focuses on street trees, which are a subset of all urban trees that possess the richest data and most consistent management across communities (Box 1). Our fitting set (used as the basis for street tree distribution extrapolation) consisted of 653 street tree databases for US communities where street tree inventory data had been collected (Fig. S3, Koch et al. 2018). In two communities (Tinley Park and, IL and Fort Wayne, IN), preventive cutting for EAB was conducted prior to the most recent inventory and was therefore accounted for within our dataset. We modelled the abundance and diameter at breast height (DBH) for trees within each genus in each community, as tree removal and replacement costs are dependent on number and diameter of trees (Aukema et al. 2011). We split trees into three DBH classes (small = 0-30cm, medium = 31-60cm, large >60cm). We first fit models for the total tree abundance of all species by DBH class, and then used these total tree models to help predict genus-specific tree abundance within each DBH class. Street tree inventory data are not always reliably reported to the species level across municipalities, and some species are so rare in street tree inventories that it would have been very difficult to develop robust species-level models, so we limited our examination to the genus level (though we note that our results did not differ qualitatively when we used more precise data; see ‘1101_severity_ by_pest_interpolate.R’ within our associated GitHub repository). Since IAFIs may not be equally impactful to all host tree species in a genus, we had to estimate the genus-level severity of each IAFI species for each IAFI-host combination. We did so by estimating the species-level breakdown of each genus based on their average relative proportions across the 653 inventoried communities within our fitting set, and assuming this distribution was representative in other projected communities.

We modelled the total abundance of street trees in a community using boosted regression trees (BRT, *gbm*.*step* within R package *dismo*, Hijmans et al. 2017, Appendix S1) relating the logarithmically-scaled total tree abundance within a DBH class to community-specific predictors, employing environmental variables from WORLDCLIM (Fick & Hijmans 2017) and community characteristics used in Koch et al. (2018), which were sourced largely from the National Land Cover Database (NLCD, Homer et al. 2015), the US Census and the American Community Survey (https://www.census.gov/data.html, Table S2).

Next, we estimated the abundance of street trees within each genus, using the same climatic and demographic factors as the total tree abundance model as well as the total tree abundance model output as predictors (Fig. S1). We considered two approaches: 1) zero-inflated Poisson generalized additive models (GAMs), or 2) a two-step BRT approach. We compared BRT and GAM models that were fitted to all genera simultaneously (general BRT/GAM models using genus-specific intercept terms) with models that were fitted to each genus separately (customized BRT/GAM models) (Fig. S1). We chose the model that produced the strongest relationship for each genus using R^2^ values that were relative to the 1:1 line (i.e., a normalized mean squared error, R^2^_MSE_, Appendix S1).

We synthesized the previous two modelling steps, intersecting IAFI spread forecasts with predicted tree distributions (using observed tree data from the fitting set where available), to create forecasts of tree exposure, which we define as the sum of predicted density of each IAFI species, multiplied by their predicted host tree abundance in each community.

### Host mortality model

We examined the impacts of the three major feeding guilds of IAFIs (Aukema et al. 2010). Foliage feeders included insects that feed on leaf or needle tissue. Sap feeders included all species that consume sap, including scale insects and gall-forming species. Borers included species that feed on phloem, cambium, or xylem. Across insect guilds, the logic from Aukema et al. (2011) appeared to hold: most species were innocuous, but a small number caused high mortality (Table S8).

We ranked the severity of a given IAFI infestation on a particular host using a scale based on observed long-term percent mortality (Table S8, defined in Potter et al. 2019). We fitted a Beta distribution to the frequency distribution of IAFI-host interactions in each of these categories using Stan (Carpenter et al. 2017), a program and language for efficient Bayesian estimation. We used the posterior mean of each severity class as the expected mortality for an IAFI-host interaction within each category.

We define the term ‘mortality debt’ as the period between an IAFI initiating damage within a community and reaching its estimated long-term (asymptotic) host mortality within that community (see Appendix S2 for more details). While we had estimates of asymptotic mortality of host trees (Potter et al. 2019), we had no information on the rate by which trees reach this plateau. Previous estimates have ranged from 5 to 100 years (Aukema et al. 2011, Pugh 2010), so we analyzed three scenarios within this range (10, 50, 100 years). To account for what is currently known about the mortality dynamics of IAFIs within each of the feeding guilds, we focused on a reasonable scenario of mortality debt across IAFI feeding guilds parameterized from several recent publications (Appendix S2) (10 years for borers, 50 years for defoliators, and 100 years for sap feeders). For simplicity, we assumed mortality increased by a constant fraction over time until reaching its maximum and levelling off. For example, in the 50-year mortality debt scenario, if an IAFI’s maximum host mortality was defined as 90%, mortality would increase by 9% at each 5-year timestep for 10 timesteps until 90% mortality had been reached.

### Management costs

As a final layer that allowed us to move from mortality estimates to cost estimates, we estimated the cost of removing and replacing dead trees. We used this cost because we believe it to be the minimum management response required, and because the extent and variability of preventive behaviour would be much harder to estimate. However, we note that this cost does not account for additional preventive cutting or any non-cutting management such as spraying or soil drenching with insecticides. We assumed that cutting was a one-time 100% effective treatment against IAFIs, or, in other words, that newly planted trees were of different species and thus not susceptible to the same IAFI species that killed the previous trees. We assumed a 2% discount rate for future damages (Aukema et al. 2011) and that infestations were independent, or, in other words, that invasion by one IAFI did not interfere with invasion by another. This is likely a justifiable assumption, as there is limited host sharing across IAFIs (but see Preisser et al. 2008), and IAFI species each infest only a small proportion of hosts at any given time interval, so there is limited potential for species interactions (Aukema et al. 2010).

We assumed the same per-tree cost estimates for cutting and replacing dead trees as in Aukema et al. (2011), where the cost of cutting increases nonlinearly with size class. If we assume that street trees are always under the jurisdiction of local governments, the cost of removal and replacement of each tree is US$450 for small trees, US$600 for medium trees, and US$1200 for large trees (these costs jump to an estimated US$600, US$800, and US$1500 for homeowners). We reported all costs incurred from 2020 to 2050 in 2019 US dollars based on a 2% discount rate relative to these baseline costs. Since these baseline per-tree management costs came from a 2011 publication, we converted them to 2019 dollars via the US Consumer Price Index (World Bank, https://data.worldbank.org). Though the tree removal and replacement cost estimates are close to those reported in other publications (Bigsby et al. 2014), these costs possess uncertainty due factors such as regional variation in labor costs, the use of external contractors, and community preferences in replacement trees. Due to the scale of our analysis, we did not attempt to quantify this additional source of uncertainty.

### Model synthesis

Once all subcomponent models had been parameterized, we synthesized the street tree estimates, IAFI spread estimates, host mortality estimates, and removal costs to produce overall cost estimates (Fig. S1). We summed the damages from 2020 to 2050 to obtain a total discounted cost for this 30-year window. We then obtained annualized costs by calculating an annuity over the 30-year time horizon using the following equation:

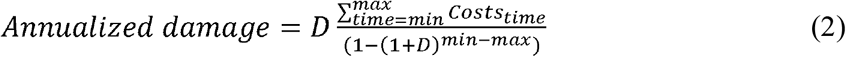

where D is the discount rate (2%).

We assessed parameter uncertainty in proportional host mortality by sampling from our posterior beta mortality distribution. We also used sensitivity analysis to explore the effect of different mortality debt scenarios, including 1) our reasonable scenario, 2) setting all guilds to 10, 50, or 100-year debts, and 3) varying each guild separately while holding the other two guilds at their reasonable scenario. While our host distribution models were based on standard modelling approaches (e.g., GAM), our Bayesian formulations underlying the mortality estimates were novel and needed to be tested theoretically, to ensure that parameters were identifiable, and reproduced the correct behavior. See Appendix S3 for details of our theoretic analyses.

### Potential impacts to non-street trees

To provide a rough estimate of non-street tree impacts (Box 1) we built a model for whole-community trees (i.e., street + non-street trees) from the dataset of 56 communities where genus-level estimates were reported, subtracted predicted street trees from this whole-community estimate, and apportioned the remaining trees into residential and non-residential trees based on their average fractions across all sites where land type breakdowns were provided (32 municipalities).

## Results

By combining these four components, we were able to forecast the sites, trees, and IAFIs associated with the greatest tree damage and associated costs across the US in the next 30 years. We found that 2.1-2.5% of all street trees will be killed in this period by IAFIs, and that this loss will cost $US 30M per year to manage. Spatially, most of the predicted damage was clustered in a 902,500km^2^ hotspot region, which is predicted to contain 95.7% of all mortality. In terms of IAFI species, EAB was associated with 85% of all costs and caused 98.8% loss of ash street trees in the hotspot region. Beyond these risks due to already-established IAFIs, we determined that future wood-boring IAFIs of Asian origin that infest maple and oak trees could lead to the greatest impact, particularly when entering from southern ports.

### Urban tree pest exposure

Total tree abundance models were predictive with some outliers (Appendix S1, Fig. S4, small trees: R^2^ = 0.78, medium trees: R^2^ = 0.58, large trees: R^2^= 0.42). Removing the outliers changed the R^2^ to 0.76 for small trees, 0.76 for medium trees, and 0.58 for large trees. Our genus-level abundance models were strong but became slightly weaker for rare genus - size class combinations (Fig. 1, overall R^2^ for all genera of small trees: R^2^= 0.93, medium trees: R^2^ = 0.93, large trees: R^2^ = 0.92). While relationships were variable across genera, the genera that were predicted most poorly did not make up a large proportion of predicted trees, and none were below R^2^ = 0.25 (Fig. S5).

**Figure 1.**
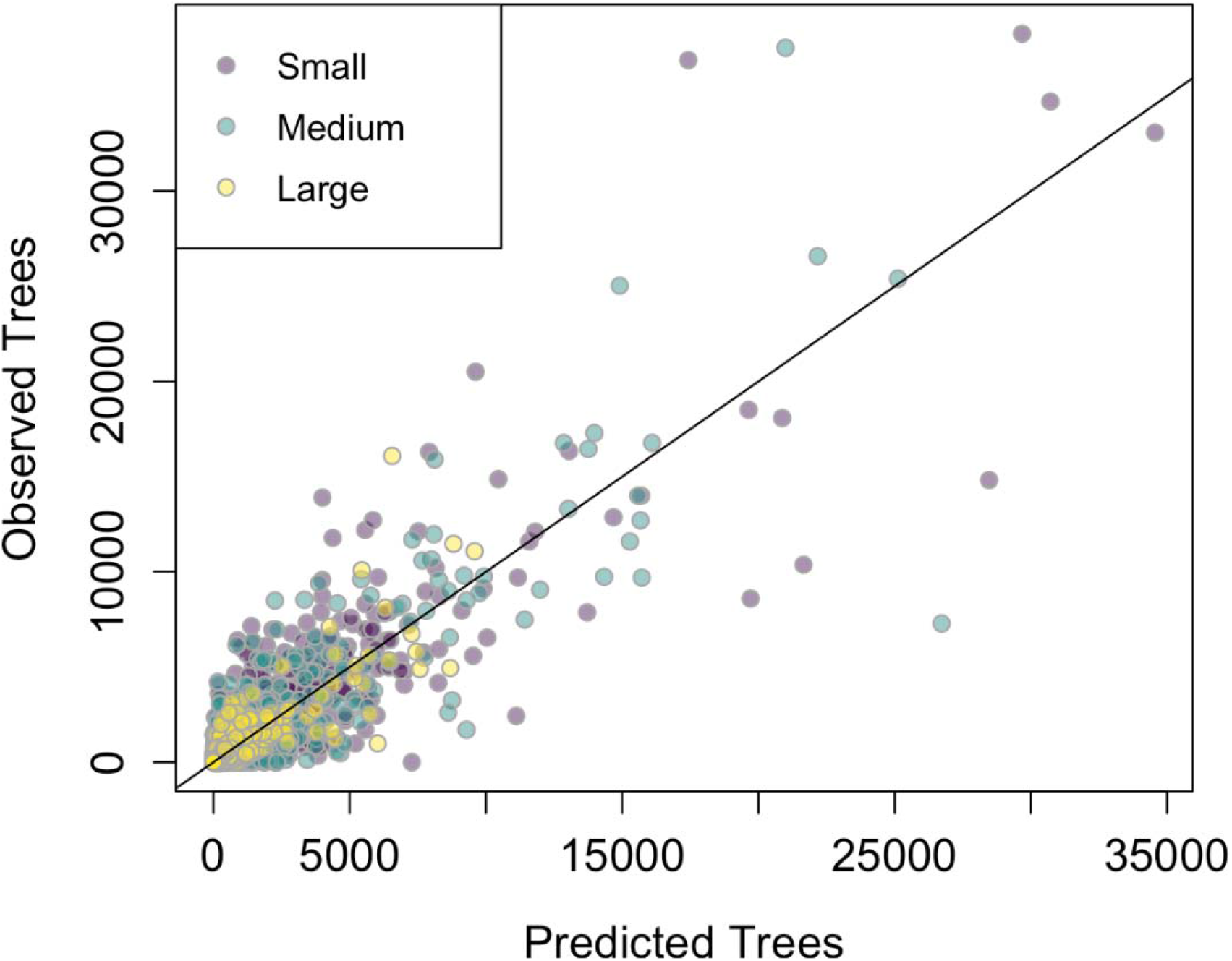
Fit of the genus-specific host tree models across all genera and size classes.

The optimal genus-level fitting approach differed across genera depending on DBH class, prevalence of genera, and whether presence/absence or tree abundance was the response variable (Table S3). Generally, rarer genera were better predicted by global BRT and GAM models, which used information from all other species, while common species were better fit by customized models (Fig. S6). According to our models, although subject to regional variation, the population of street trees is mostly made up of maple (*Acer*) and oak *(Quercus)*, with substantial ash (*Fraxinus*, Fig. S7).

Predicted street tree exposure (measured as the number of predicted susceptible trees in Fig. 2a * IAFI relative propagule pressure in Fig. 2b, Hudgins et al. 2020) across all tree types from 2020 to 2050 was generally high in the eastern US, and only sporadically high across the western US (Fig. 2c). Predicted street tree exposure was highest among maples (*Acer* spp., 25.6M predicted exposed trees), oaks (*Quercus* spp., 5.9M), and pines (*Pinus* spp. 3.4M). This latter genus was largely confined to the Southwest but had high exposure to scale insects. The greatest number of trees were predicted to become exposed to San Jose scale (*Quadraspidiotus perniciosus*, 7.3M), Japanese beetle (*Popillia japonica*, 6.7M), and calico scale (*Eulecanium cerasorum*, 6.4M). Among residential and community trees, exposure was greatest among maples, oaks, and *Prunus* spp. (1.7B,1.1B, 707M, respectively), and the most frequently predicted IAFI encounters were with the same three species (Japanese beetle, San Jose scale, and calico scale).

**Figure 2.**
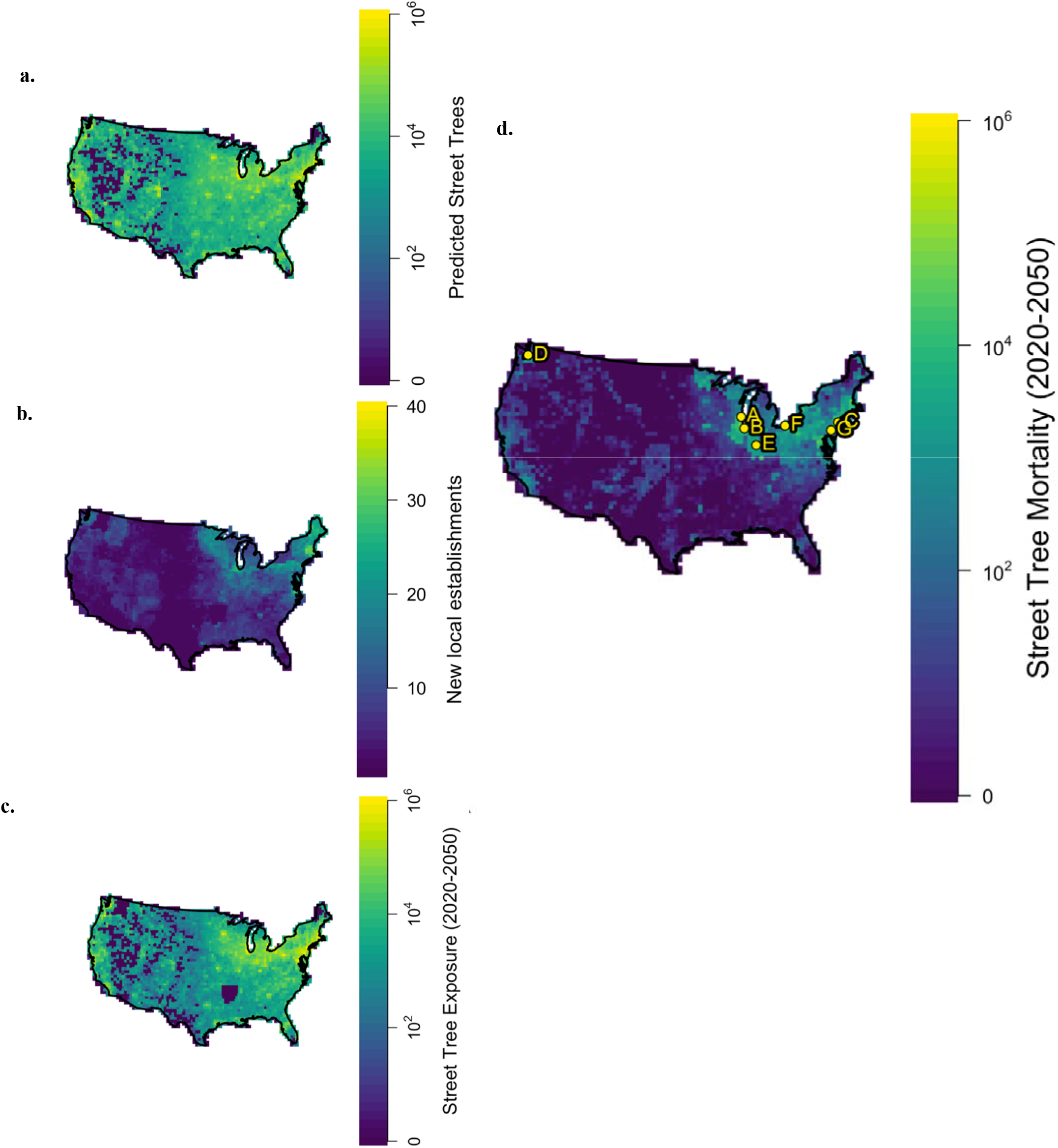
Model outputs for the first three subcomponent models, including **a**. predicted street tree abundance, **b**. predicted newly invaded sites of existing IAFIs, **c**. predicted street tree exposure levels (number of focal host tree + IAFI interactions) from 2020 to 2050, and finally **d**. Predicted total tree mortality from 2020 to 2050 in the reasonable mortality debt scenario across space. The top seven most impacted cities or groups of nearby cities are shown in terms of total tree mortality 2020 to 2050 (A = Milwaukee, WI; B = Chicago/Aurora/Naperville/Arlington Heights, IL; C = New York, NY; D = Seattle, WA; E = Indianapolis, IN; F = Cleveland, OH; G = Philadelphia, PA).

### Host tree mortality

The best-fitting mortality model indicated that most IAFIs fall in the low severity groups. Within all severity groups, the majority of IAFIs were at the low end of severity (Fig. 3, full results in Appendix S2). In our reasonable mortality debt scenario (i.e., 10-year scenario for borers, 50-year scenario for defoliators, 100-year scenario for sap feeders, we estimated a mortality level of 0.7-2.5% above expected background mortality of street trees by 2050, where our reasonable scenario fell on the higher end of this range (Table 1). Predicted street tree death varied by a factor of four based on the mortality debt scenario, with longer debts leading to lower total mortality between now and 2050 (Table 1). This was because in longer mortality debt scenarios, trees experience mortality for 50 or 100 years after pest establishment, compared to only 10 years in our shorter mortality debt scenario. Therefore, for our highest impact IAFI (EAB), which has only established recently, much of its mortality would be incurred in the years after 2050 in the 50 year and 100 year mortality debt scenarios. Mortality estimates are most sensitive to changes in the impact of wood boring species, as demonstrated by the sensitivity of mortality estimates to their mortality debt scenarios (“Vary Borers” row, Table 1). We also found that longer mortality debts led to a smoother cost curve, or costs that do not vary much due to more consistent host mortality rates (Fig. 4).

**Figure 3.**
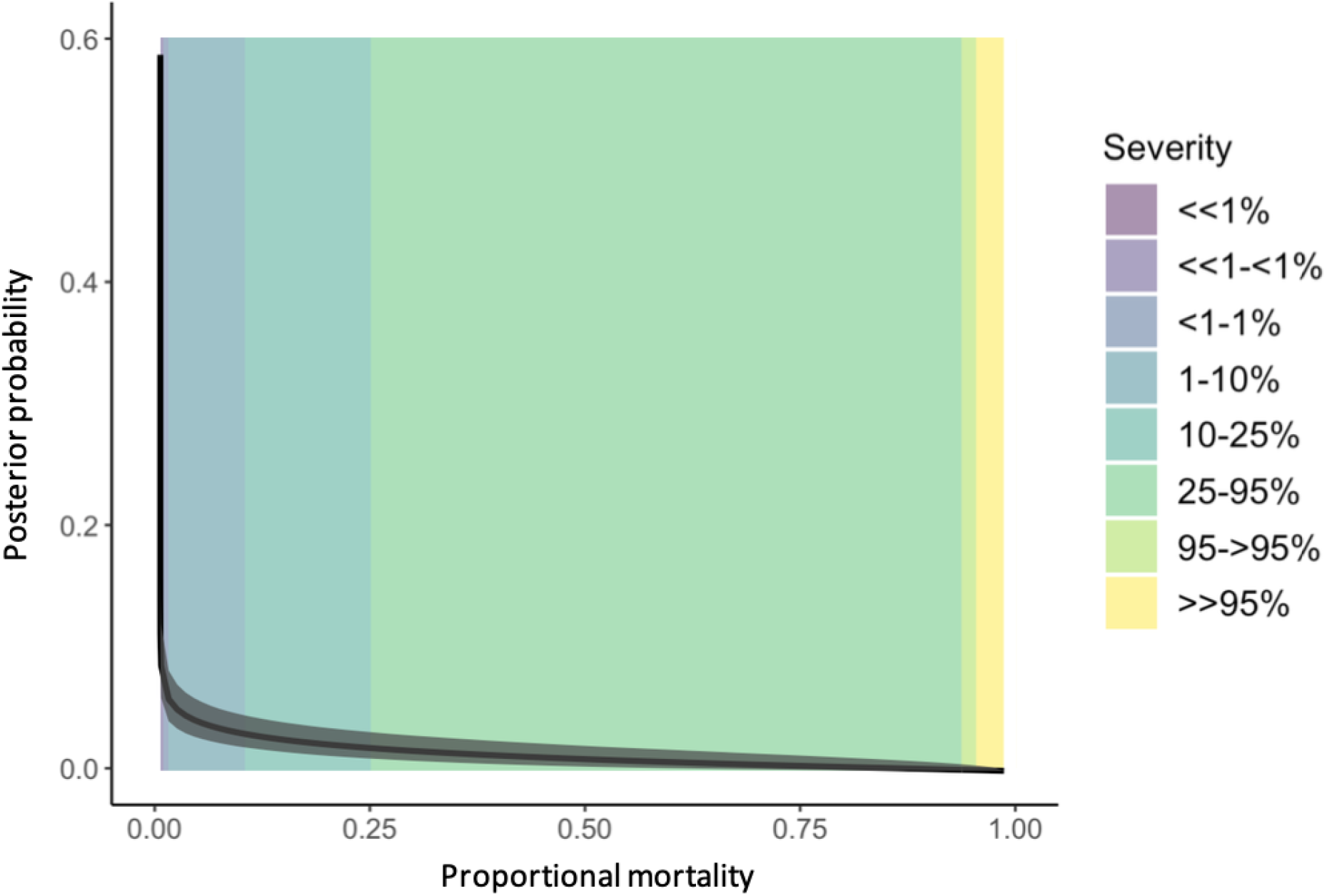
Posterior distribution for the beta model of host mortality due to IAFIs within each severity category. 95% Bayesian credible intervals are shown in grey, and the posterior median is shown in black. Colored bins represent severity categories extended from Potter et al. (2019).

**Table 1.** Predicted annualized cost (in 2019 US dollars) and tree mortality across invasion scenarios from 2020 to 2050 across all 57 IAFI species. “Reasonable” indicates the scenario with mortality debt durations determined by recent publications (Appendix S2). “Vary” scenarios hold all guilds but the focal guild constant at their reasonable scenario, and “All” fix all three guilds at a given mortality debt duration. Mean mortality for reasonable scenario = 2.3%, 1.38M trees, US$ 30M annualized (US$ 679M over the next 30 years).

| Mortality Debt Scenario | Annualized Cost (US\$ millions) | | Tree Mortality (Millions) | | Percent Mortality | |
| --- | --- | --- | --- | --- | --- | --- |
|  | lower 95% CI | upper 95% CI | lower 95% CI | upper 95% CI | lower 95% CI | upper 95% CI |
| Reasonable | 28.5 | 33.2 | 1.29 | 1.54 | 2.1% | 2.5% |
| Vary Borers | 10.1 | 32.1 | 0.45 | 1.45 | 0.7% | 2.4% |
| Vary Defoliators | 28.1 | 32.6 | 1.28 | 1.48 | 2.1% | 2.4% |
| Vary Sap-feeders | 28.5 | 32.5 | 1.30 | 1.47 | 2.1% | 2.4% |
| All 10 | 27.8 | 30.4 | 1.27 | 1.39 | 2.1% | 2.3% |
| All 50 | 18.5 | 22.3 | 0.84 | 1.00 | 1.4% | 1.7% |
| All 100 | 9.77 | 13.5 | 0.44 | 0.60 | 0.7% | 1.0% |

**Figure 4.**
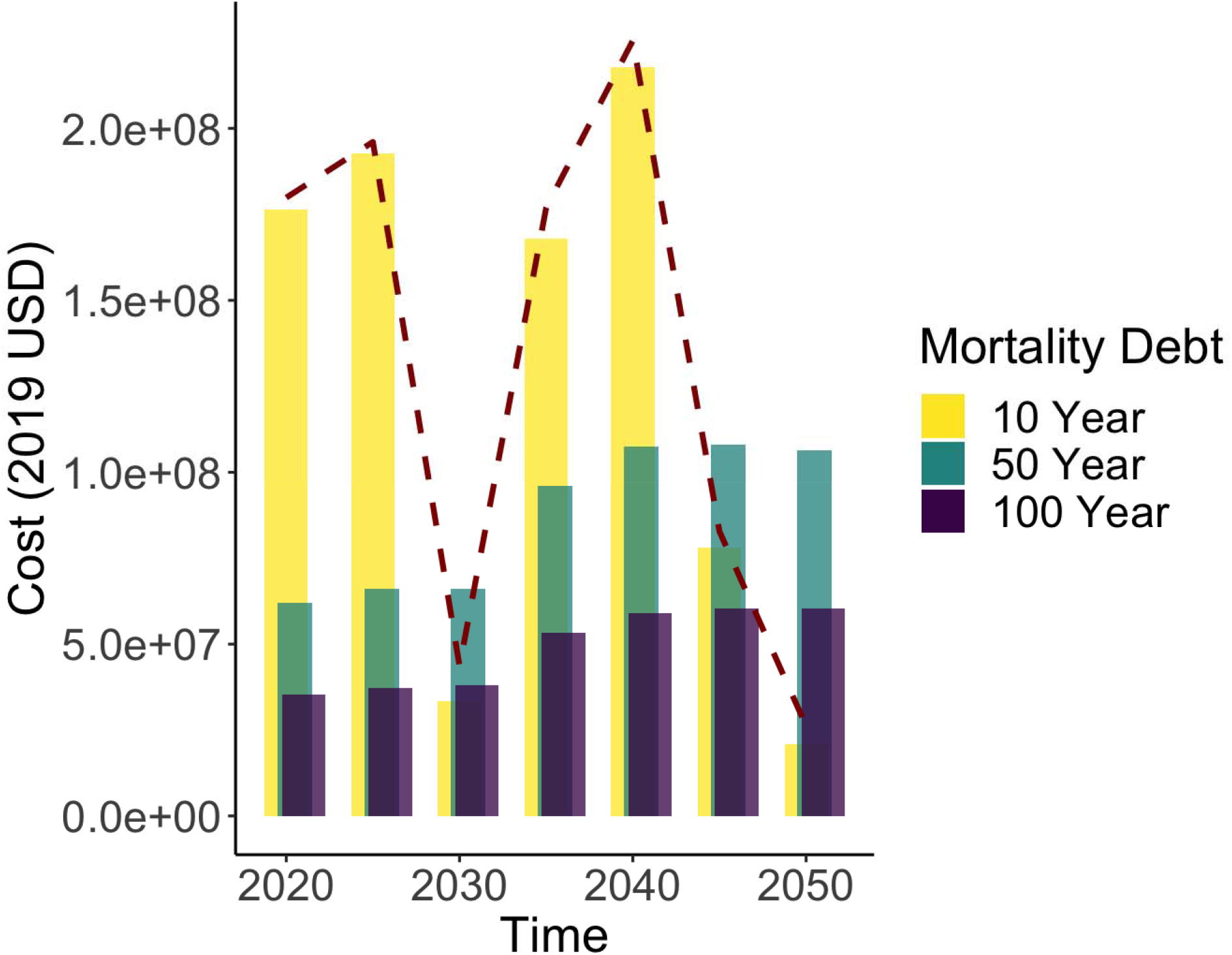
Depiction of the influence of mortality debt on temporal cost patterns. Predicted costs 2020 to 2050 for the 10 year (yellow), 50 year (teal), and 100 year (purple) mortality debt scenarios with a 10 year initial invasion lag. The reasonable scenario predictions are shown as a dashed red line. Costs are presented in 5-year increments in accordance with the timestep length within our spread model.

Spatially, future damages will be primarily borne in the Northeast and Midwest, driven by EAB spread (Fig. 2d). We predict that EAB will reach asymptotic mortality in 6747 new cities, which means that 98.98% of its preferred ash hosts will die. Thus, the mortality is predicted to be concentrated in a 902,500km^2^ zone encompassing many major Midwestern and Northeastern cities (Fig. S10). This mortality is also predicted to result in a 98.8% loss of all ash street trees within this zone. Examining the back-casts available within our model results, we see that over 230,000 ash street trees are predicted to have died before 2020. On the other extreme, when we use our model to forecast even longer into the future, there are a further 69 cities where EAB is predicted to reach asymptotic mortality within 10 years of 2050 (i.e., 98.8% ash mortality by 2060). Furthermore, at-risk ash trees are unequally distributed. We projected the highest risk close to the leading edge of present-day EAB distributions, particularly in areas predicted to have high ash densities. The top “mortality hotspot cities”, where projected additional mortality is in the range of 5,000-25,000 street trees, include Milwaukee, WI; the Chicago Area (Chicago/Aurora/Naperville/Arlington Heights, IL); Cleveland, OH; and Indianapolis, IN (Fig. 2d). Cities predicted to have high mortality outside of the Midwest include New York, NY; Philadelphia, PA; and Seattle, WA – communities with high numbers of street trees and high human population densities, which attract EAB propagules within our spread model. The states most impacted by street tree mortality match these patterns, where the highest mortality is predicted for Illinois, New York, and Wisconsin.

### Cost estimates

We estimated annualized street tree costs across all guilds to be between US$29-33M per year in our reasonable scenario (mean = $30M, Table 1, Fig. S11). Roughly 90% of all costs across the entire US were due to EAB-induced ash mortality. The total cost associated with street tree mortality in the top ten hotspot cities was estimated at $50M from 2020 to 2050, with $13M in Milwaukee, WI alone.

The ranking of feeding guild severity was relatively robust across mortality debt scenarios, despite the potential for differences due to the interaction of IAFI-specific spread and mortality debt dynamics. Costs were higher for longer mortality debt scenarios for borers, peaked at intermediate debt for defoliators, and peaked at the longest debt for sap feeders. These patterns were due to the relative rates of historical and contemporary range expansion of more impactful IAFIs (i.e., high impact borers have more rapid recent range expansion, while contemporary high impact defoliator expansion is slow compared to 50 years ago). Borers were predicted to be the most damaging feeding guild ($8M-28M mean annualized street tree damages across scenarios), and EAB was consistently the top threat. Defoliators were predicted to be the second most damaging feeding guild in the next 30 years (means = $0.8M-$1.4M), despite having more widespread hosts than wood borers, due to lower asymptotic mortality levels. Defoliators had a 1-2 orders of magnitude lower cost than wood-boring species, but again showed consistency in which species were the top threats within the guild. Consistent with previous work in Aukema et al. (2011), *Lymantria dispar dispar* (LDD) moth had the highest cost of all defoliators, followed by Japanese beetle and cherry bark tortrix (*Enarmonia formosana*). The sap-feeding group accrued the lowest costs in the next 30 years due to their lower asymptotic mortality and rarer street tree hosts (mean = $0.2M-1.1M). Hemlock woolly adelgid (*Adelges tsugae*) was the highest impact sap feeder, followed by oystershell scale (*Lepidosaphes ulmi*) and elongate hemlock scale (*Fiorinia externa*). Total costs were sensitive only to borer mortality debt scenario specification (Table 1).

### Potential impacts to non-street trees

Mean additional mortality (i.e., above background rates) for residential and non-residential community trees in the reasonable scenario was 1.0% (13.3M residential and 72.1M non-residential trees, Table S11). While recognizing that non-street tree management will likely be more variable, to provide a rough estimate, we assumed that non-street trees would be managed in the same way as street trees (i.e., removal and replacement of dead trees). In this scenario, additional mortality would incur an estimated annualized cost of $1.5B for non-residential trees and $356M for residential trees. Further, a disproportionate amount of the total damages (91% of the mortality to residential and non-residential community trees) is expected to be felt in the hotspot zone, with 12.1 million residential and 65.9 million non-residential community trees expected to be killed. This is most certainly a high-end estimate for non-residential trees, because many of these trees are at a very low priority for removal, such as those in open spaces and in urban forests. Given the relatively limited data, and the difference in potential management behaviour for these trees, we caution against overinterpretation of these results.

### Novel IAFI risk forecast

Our framework allowed us to identify the factors leading to the greatest impacts for IAFIs already known to have established in the United States. We were able to identify the most common urban host trees, the sites facing the greatest future IAFI propagule pressure, and the IAFI-host combinations with the greatest mortality. However, this approach can also be synthesized with IAFI entry scenarios to understand potential impacts of novel invasive IAFIs. To illustrate the utility of this framework for prediction, we have provided a checklist of risk factors in Table S12 and future spread simulations in Table S13 and Fig. S12. We show that entry via a southern port (e.g., the Port of South Louisiana) would lead to the greatest number of exposed trees. Further, an EAB-like borer of oak and maple trees could kill 6.1 million street trees and cost $4.9B over the next 30 years.

## Discussion

While previous analyses have indicated that urban trees are associated with the largest share of economic damages due to IAFIs (Aukema et al. 2011, Paap et al. 2017, Kovacs et al. 2010), until recently, data did not exist on the urban distribution of host trees (Koch et al. 2018), the spread of IAFIs (Hudgins et al. 2017,2020), or the mortality risk for hosts due to different IAFIs (Potter et al. 2019). With these new models, it is now possible to forecast where and when IAFIs will have the most damages across the US. Our analysis suggests an overall additional mortality of between 2.1-2.5% of all street trees, amounting to $US 30M per year in management costs. However, the most useful element was our ability to forecast hotspots of future forest IAFI damages, including a 902,500km^2^ region that we expect to experience 95.7% of all mortality, in large part due to a 98.8% loss of its ash street trees due to EAB. This type of forecasting has been highlighted as a crucial step in prioritizing management funds (McGeoch et al. 2016). These data can be used by municipal pest managers to anticipate future costs and may help motivate improved spread control programs that aim to identify the potential source locations of future invasions and mitigate the worst anticipated impacts (complete forecast available at http://github.com/emmajhudgins/UStreedamage). As an example, cities could use these forecasts to determine their need for the deployment of trapping or other early-detection efforts. Cities expected to face large losses in the coming years are largely concentrated in the US Midwest, where the current EAB invasion is most severe, but also include areas thousands of kilometers to the west, such as Seattle, WA. These communities could benefit more from increasing their surveillance, while cities outside future hotspot regions may require less investment in these tools.

Beyond present IAFI risks, our integrated model can also act as a risk assessment tool for street tree mortality caused by novel IAFIs (Table S12-S13, Fig. S12). Although ash trees are assured to be dramatically affected by EAB over the next few decades, our models suggest oak and maple to be the most common street tree genera nationwide. Further, while ash species are being substituted with less susceptible tree species, maples and oaks continue to be widely planted within our street tree inventories. Therefore, IAFIs with host species in these genera should be of heightened concern. Secondly, the timescale and magnitude of the impacts of wood borers (see also Aukema et al. 2011) make them the highest risk to street trees. We integrated these two pieces of information with information on major ports of entry within the US (American Association of Port Authorities 2015, http://aapa.com/), as well as our general model of IAFI spread (Hudgins et al. 2020), to forecast the extent of exposed maple and oak street trees from 2020-2050 (Fig. S12, Table S13). Our analyses show that entry via a southern port would lead to the greatest number of exposed trees. Moreover, larger trade volumes between the US and Asia compared to other regions (Sardain et al. 2019) suggest Asian natives will be the most likely future established IAFIs (see also Koch et al. 2011). One potential candidate species fitting these criteria is citrus longhorned beetle (*Anoplophora chinensis*), which is an Asian wood borer possessing many potential US host species, including ash, maple and oak (Haack et al. 2010). This species was discovered in a nursery in Tukwila, WA in 2001 and rapidly eradicated, and so far, no establishments have been found (Haack et al. 2010). The lack of more thorough regulation of live plant imports and strict implementation of current wood treatment protocols such as ISPM15 (Lovett et al. 2016) increase the susceptibility of the US to invasion and subsequent spread of this species and other potentially high-risk borers. Note that our risk forecast is for post-establishment spread only, and does not attempt to model the risk of entry. We note that while *Anoplophora* species may have lower climatic suitability in the southern US (see Byeon et al. 2021), propagule pressure can lessen the role of climatic limitations in invasion processes, especially those fitted based on existing invasion data (Bradie & Leung 2015).

Our impact estimates vary substantially based on dynamics of host mortality following initial IAFI invasion, especially because of variability in the duration and functional form of mortality debt. As it happens, we are most confident in our mortality debt specification for the guild (borers) and species (EAB) whose impacts on total community costs are most sensitive to mortality debt. Several publications have demonstrated near-complete decimation of ash stands in the decade following EAB infestation (Kovacs et al. 2010, Knight et al. 2013, Fei et al. 2019). Furthermore, since total tree mortality is asymptotically equivalent across all mortality debt regimes, if other feeding guilds possessed 10-year mortality debt regimes, we should have been able to detect a rapid die-off of their hosts as they spread, similarly to what we found for EAB (albeit scaled by their maximum mortality rates). This is not the case in the literature (Fei et al. 2019).

With our integrated model, we also estimated economic damages, which updates the decade old Aukema et al. (2011) using recent advances (Koch et al. 2018, Potter et al 2019). We estimate annualized total costs to urban trees to be somewhat lower than those in Aukema et al. (2011). Namely, we estimate $2B in our cases versus the $2.0B in total “Local Government expenditures” and 1.1B in “Household Expenditures” reported in Aukema et al. (2011). Our lower overall estimate is likely because of a lower rate of predicted ash exposure to EAB (i.e., lower predicted ash abundance in areas of predicted EAB spread) in non-residential areas. However, the methodologies used in this previous analysis are so different that a direct comparison of the subcomponents of the expenditures is not possible.

While EAB drove the patterns of impact across the country, it is noteworthy that the impact dynamics of other feeding guilds followed a qualitatively similar pattern, with the highest impacts in the northeastern United States (Fig. S13). LDD moth drove defoliator costs, which appear somewhat more concentrated in the northern US than the other two guilds, although there is also a hotspot around Seattle, WA. The sap-feeding group’s costs were driven by hemlock woolly adelgid (*Adelges tsugae*) and were concentrated in the southwestern US compared to defoliators, particularly in California.

We predict that the majority of communities containing ash trees will not have reached their maximal EAB-induced ash mortality by 2060, because of lower densities of forest ash beyond our forecasted invasion extent, thus limiting exposure. Spatially, our results show lower threat in the western US. This pattern is consistent with previous findings (Lovett et al. 2016) and can be explained by the high impacts of EAB, LDD moth, and hemlock woolly adelgid, whose distributions are projected to remain concentrated further east in the short term. However, some of the highest-impact non-native pathogens have emerged in the western US and were not captured in this analysis (Kinloch Jr. 2003, Rizzo & Garbelotto 2003). Western regions could also see high future risks due to the polyphagous shot hole borer (*Euwallacea whitfordiodendrus)* and its insect-disease complex with fusarium fungus (*Fusarium spp*.) (Coleman et al. 2019). This complex has already established in California and has maple and oak trees among its many hosts.

While the substantial advances that emerged recently allowed us to develop a more fully integrated model, we also identified data deficiencies which require additional research. A relative quantification of additional sources of uncertainty is provided in Table S14. Furthermore, this cost estimate only examines the cutting of dead trees. The analysis fails to account for preventive cutting prior to EAB arrival, to fully examine non-street tree management, and to assess the impacts of IAFIs that have not yet established in the United States. Moreover, our analysis assumes a complete identification of ‘high impact IAFIs’. Some presently established IAFIs may not yet have been identified as ‘high impact’, either due to lags in their impact and/or lags in the detection of this impact (Coutts et al. 2018), but may become ‘high impact’ before 2050. Beyond the 57 IAFIs examined, novel invaders may establish between now and 2050 and begin to cause additional impacts. Finally, our analysis does not capture the myriad other impacts of IAFIs, including the substantial ecosystem services losses they are known to cause (Hill et al. 2019).

We have shown that the suite of known IAFIs have the potential to kill roughly a hundred million additional urban trees in the US in the next 30 years. While these numbers themselves are striking, reporting only a country-level impact estimate without IAFI species, tree, and community-level resolution does little to inform management prioritizations. Here, we were able to identify specific urban centers, IAFI species, and host tree genera associated with the vast majority of these impacts. We predict that 90% of all street tree mortality within the next 30 years will be EAB-induced ash mortality, and that ∼95% of all street tree mortality will be concentrated in less than 25% of all communities. These estimates illustrate the gravity of IAFI infestations for communities in the path of high impact invaders that are rich in susceptible hosts. Moreover, we were able to use this framework to identify a checklist of biotic and spatiotemporal risk factors for future high-impact street tree IAFIs.

## Acknowledgments

EJH would like to thank her PhD supervisory committee members T. Jonathan Davies and Patrick M. A. James for their invaluable comments, as well as the thoughtful comments and questions from thesis external examiner Dominique Gravel and colleague Andrew Liebhold. EJH also acknowledges the continual support and feedback from lab members Dat Nguyen, Abbie Gail Jones, Charlotte Steeves, Shriram Varadarajan, and Lidia Della Venezia. This work was supported by a NSERC CGS-D awarded to EJH. This research was funded in part by the US Department of Agriculture, Forest Service, including through Cost Share Agreements 17-CS-11330110-025, 18-CS-11330110-026, and 19-CS-11330110-027 between the Forest Service and North Carolina State University.

## Boxes

### Box 1.

#### Urban tree populations modelled in this analysis.

**Urban trees**: The complete population of trees found within the limits of a community.

**Street trees:** The subset of the urban tree population planted alongside roads (including on residential properties) and usually managed by local municipalities.

**Non-street trees:** The subset of the urban tree population not planted near roads (including residential and non-residential trees)

**Residential trees:** A subset of non-street trees planted on residential properties. These trees most often are managed by individual property owners.

**Non-residential trees:** A subset of non-street trees not planted on residential property, including trees in parks and other municipal properties, cemeteries, and undeveloped property, as well as trees on commercial/industrial property.

## Notes

**Conflict of interest:** The authors have no conflicts of interest to declare.

https://github.com/emmajhudgins/UStreedamage

### Competing Interest Statement

The authors have declared no competing interest.

### Summary of Updates

This is the revised version of the manuscript resubmitted to Journal of Applied Ecology following minor revisions. Two typos were corrected and one sentence was reworded for clarity.

