## Appendices for "Hotspots of pest-induced US urban tree death, 2020-2050"

**This PDF file includes:**

Supplementary text

Figures S1 to S12

Tables S1 to S14

Legend for Dataset S1

SI References

**Other supplementary materials for this manuscript include the following:**

Dataset S1

Supplementary Information Text

Appendix S1. Host distribution model: detailed methods and results.

Extended methods

We hypothesized that the age and wealth of a community would influence the types and sizes of trees planted there. In our model, median home value and mean year of construction (at the block-group level) as well as median household income (at the county level) were used as proxies of the age of the urban tree community and the community budget for street trees. We also tested the use of Poisson GAM models, but high levels of concurvity (the GAM equivalent of multicollinearity, [1] amongst predictors and lower predictive performance indicated Poisson GAMs were an inferior modelling structure for estimating total abundance.

For BRT, we modeled tree presence/absence, followed by tree abundance given presence (using logarithmically-scaled tree abundance and back-transforming when predicting), and then combined the two models. The number of trees of genus *i* in size class *j* at a particular site *k* was:

$\boldsymbol{trees}_{\boldsymbol{i,j,k}}\boldsymbol{=}{\boldsymbol{c}_{\boldsymbol{i,j,k}}\boldsymbol{*pred}}_{\boldsymbol{exist,i,j,k}}\boldsymbol{*}\boldsymbol{pred}_{\boldsymbol{number,i,j,k}}$	(1)

$\boldsymbol{c}_{\boldsymbol{i,j,k}}\boldsymbol{=}\frac{\boldsymbol{1}}{\sum_{\boldsymbol{k}} \boldsymbol{(}\boldsymbol{pred}_{\boldsymbol{exist,i,j,k}}\boldsymbol{*}\boldsymbol{pred}_{\boldsymbol{number,i,j,k}}\boldsymbol{)/}\sum_{\boldsymbol{k}} \boldsymbol{obs}_{\boldsymbol{number,i,j,k}}}$	(2)

This process is similar to a zero-inflated Poisson (ziP) model [2] but does not link the parameters of the binary and continuous components of the model, instead fitting them separately. Because our BRT approach was built from two independent parts, we needed to add a rescaling step so that the output summed to the observed counts (eqn. 2), as occurs for ziP models by default [2]. We removed all highly correlated variables (r > 0.8) prior to fitting, and refit GAMs until maximum estimated worst-case concurvity using three-knot smoother functions was below 0.8 (*concurvity* function within *mgcv,*[3]).

We compared BRT and GAM models that were fit to all genera simultaneously (general BRT/GAM models using genus-specific intercept terms) with models that were fit to each genus separately (customized BRT/GAM models) (Fig. S1). Predictive power could be higher when modelling all genera together if the genera respond similarly to predictors, while power could be higher for individually fitted genera where environmental and community characteristic relationships are idiosyncratic and where the sample is sufficiently large.

We chose the model that produced the strongest relationship for each genus using R^2^ values that were relative to the 1:1 line (i.e, a normalized mean squared error, R^2^_MSE_).

R^2^_MSE_ more correctly measures deviations between observations (y) and predictions ($\hat{\boldsymbol{y}}\boldsymbol{)}$ than conventional R^2^.

$\boldsymbol{R}_{\boldsymbol{MSE}}^{\boldsymbol{2}}\boldsymbol{=1-}\frac{\sum{\boldsymbol{(y-}\hat{\boldsymbol{y}}\boldsymbol{)}}^{\boldsymbol{2}}}{\sum{\boldsymbol{(y-}\bar{\boldsymbol{y}}\boldsymbol{)}}^{\boldsymbol{2}}}$	(3)

We removed New York, NY from the fitting set as it was likely to be a high leverage observation and could have significantly changed the resulting models due to it possessing a markedly different street tree genus composition from all other communities. Both the GAM and BRT models were fitted using their built-in cross-validation algorithms for parameter estimation, and can therefore tolerate occasional outliers with minimal effect on their parameter estimates (though we have less evidence that other outliers would have changed model parameters for cities other than New York, NY). Given the higher data requirements of GAMs (i.e. all parameters must be fit simultaneously, rather than BRT, which can fit subsets of predictors to each tree, [4]), genus-specific GAMs were not considered when data were insufficient (i.e., when only a few cities contained that genus). For each genus, we used the best-fitting model to predict urban tree distributions throughout the contiguous US. We used the observed number of trees rather than model predictions in cities where these data were available. Alaska and Hawaii were removed to match the spatial extent of IAFI spread predictions, and because urban tree genus composition is likely quite different in these areas compared to the contiguous US.

Results - Total tree abundance models

Total tree abundance models were moderately predictive with some outliers (Fig. S4, fitted R^2^ for small trees = 0.78, medium trees = 0.58 large trees= 0.42). Removing the outliers changed the fitted R^2^ to 0.76 for small trees, 0.76 for medium trees, and 0.58 for large trees. Both demographic and environmental predictors were important in predicting total trees, with human population size, ecological province, and community area emerging as top predictors across size classes, mean year of home construction emerging as predictive for medium and large trees and community canopy cover for large trees (Table S3).

We found that both demographic and environmental predictors were important in determining street tree distributions, including human population size, ecological province, community area, mean year of home construction and community canopy. In comparison, [5] also found canopy cover and area to be highly predictive of total community basal area in their models. However, these authors did not find housing or human population variables to be predictive, but instead found latitude and longitude to be predictive. Given that their analysis was restricted to three genera in the Eastern and Central US, it may have been easier to capture larger climatic patterns in the distribution of these trees in these areas, and latitude and longitude act might have served as surrogates for other factors in their restricted area of analysis.

Results - Genus-specific models

The total tree abundance models were used as inputs to the genus-level models, which were then used to predict abundances for each host tree genus. Our genus-level model fits were strong, but became slightly weaker for rare genus - size class combinations (Fig. 1, overall R^2^ for all genera of small trees = 0.93, medium trees = 0.93, large trees = 0.92). Within each genus, the best combined model fits were generally strong (Supplementary Dataset 1, mean R^2^=0.76, σ=0.16 for small trees, mean R^2^=0.78, σ=0.14 for medium trees, mean R^2^=0.78, σ=0.17 for large trees). While relationships were variable across genera, the genera that were fit most poorly did not make up a large proportion of predicted trees, and none were below R^2^ = 0.25 (Fig. S5, Table S4). Some genera were very rare within our inventoried communities at a given size class, and therefore had insufficient data to fit customized genus-level models (but could still be fit in global BRT and GAM models). In these cases, R^2^ is reported as NA in Supplementary Dataset 1.

The optimal genus-level method (global BRT, global GAM, customized BRT, or customized GAM, Fig. S5) differed across genera depending on diameter class, prevalence of genera, and whether presence/absence or tree abundance was the response variable (Tables S4-6). Across diameter classes, a general GAM for presence was most frequently selected when the number of predicted trees and the number of sites with trees present for that genus were high, indicating that the presence of common genera is driven by the same environmental and social variables. When predicting tree abundance, given presence, more common genera tended to be fit better by customized BRT or GAM models, indicating more idiosyncratic relationships to predictor variables at the level of tree quantity for ubiquitous genera. Where the number of total predicted trees and sites was not as high, small and medium size classes tended to be predicted best by a global BRT for both tree presence and abundance, while large trees tended to be fit best by a customized GAM for presence, followed by a global BRT for tree abundance. It appears that even in cases of data scarcity, genus-specific predictors of presence/absence can be detected for some rare genera. However, for tree abundance, rarer genera are better fit by general models, potentially due to their increased statistical power (Fig. S6).

According to our models, across the US, the population of small street trees is mostly made up of *Acer* and *Quercus*, with substantial *Fraxinus* (Fig. S7). Medium trees are again mostly *Acer*, but with a large number of *Eucalyptus* and *Fraxinus*. The largest street trees are the most evenly split across genera, but dominated by *Acer* and *Quercus*. When we compared our fitting set (653 communities) to our extrapolations of tree distributions across all communities in the US (~30 000 communities), small tree predictions remained relatively close to the inventoried distribution. Conversely, medium trees were predicted to have a much higher proportion of *Eucalyptus* than in the fitting set (we note these were predicted almost entirely in California). Similarly, we predicted a much smaller proportion of medium-sized *Quercus* than in the fitting set, and a much smaller proportion of large-sized *Ulmus*. These discrepancies are likely due to tree community differences in the southern US, where we had very few tree inventories, compared to the northern communities that comprised the majority of our data.

**Appendix S2.** Mortality debt – further details

EAB is estimated to kill the majority of its susceptible hosts in the first decade following infestation (Kovacs et al. 2014), while maximum mortality is estimated to take closer to 100 years for hemlock woolly adelgid (Aukema et al. 2011), so we used the 10 and 100-year scenarios for borers and sap-feeders, respectively. A recent publication examining mortality rates in forested areas suggested that *Lymantria dispar dispar* (LDD moth) has a mortality rate intermediate between borers and sap-feeders, so we set defoliators at 50-years (20). Once an IAFI was predicted to infest an area, we imposed a 10-year initial lag phase between IAFI arrival at a site and the initial onset of damage (Hochberg & Weiss 2001, Liebhold & Tobin 2008) and then began increasing the host mortality following our mortality debt scenario to the asymptotic level (defined by the host mortality model).

The joint impact of maximum mortality and mortality debt is best illustrated by a series of examples. Estimates of street tree natural mortality range around 2.4-2.6% per year (Hilbert et al. 2019). Within a 30-year window, this would amount to roughly 53% natural street tree mortality. Our model assumes that if IAFI enters site at the beginning of this window (2020), it first undergoes a 10-year time lag, and can then cause mortality in the final 20 years. The maximum level of mortality induced by a borer (EAB on several *Fraxinus* spp., Category H = 98.98%), would result in 98.98% additional mortality (mortality of remaining the trees that survived natural mortality) at the end of a 30-year window. This level of mortality would be clearly detectable above natural street tree mortality. Hemlock woolly adelgid has a similar maximum mortality to EAB (Category H on *Tsuga* spp*.*), but we have assumed that sap-feeder mortality takes 100 years to reach asymptotic levels. As such, by the end of a 30-year window, only (98.98%/100)*20 years = 19.80% of additional host trees would be killed. While defoliators have shorter mortality debts, they tend to cause lower mortality, making their impacts the least detectable above background mortality. For defoliators, the IAFI with the greatest damage on any host is the larch casebearer, (*Coleophora laricella* on *Larix laricina*, category E = 16.46%). Given a 50-year mortality debt for defoliators, the maximum mortality above background rates by 2050 is (16.46% / 50) * 20 years = 6.58%. While these estimates are much lower, many host trees of sap feeders and defoliators are very common, and this mortality could very well be inflating the perceived background mortality rates of these host trees measured in (Hilbert et al. 2019).

Appendix S3. Summary of Bayesian theoretic analyses

We ranked the severity of a given pest infestation on a particular host using a scale based on observed long-term percent mortality (defined in [6], Table S8). We added two additional categories to this scale to represent pest species missing from their database that are still considered pests on a particular host within [7]. The lowest-impact pest-host combinations were those featuring pests reported as ‘low impact’ in [7].These accounted for most known combinations. The second lowest category featured ‘intermediate impact’ pest species from [7] that did not appear as threats to a given known host in [6]. We assumed that, were these species quantified by [2], their associated severities would be lower than the lowest category within the authors’ ranking scheme. All other pest-host combinations were assigned to the same categories as in [2]. Pest frequency within severity categories was normalized across the sum of their known hosts so that each pest had equal impact on the frequency distribution (i.e., frequency summed to 1 for each pest). For instance, if a pest had 3 hosts, and had severities of 3, 5, and 9 on each host, we would give them a frequency of 1/3 under each bin. Additionally, we fit the upper limit of the two lowest mortality categories and the lower limit of the highest category, as these categories did not have quantified bounds, but could be ranked relative to others. We chose to fit a Beta distribution because proportional mortality ranged between 0 and 1.

To ensure our resulting mortality and cost curves were identifiable and had unbiased parameters, we tested their ability to fit to simulated mortality and cost data using theoretical simulations. In our theoretic analyses, we tested a series of parameter values for five curve families (the beta family for the mortality curves and gamma, lognormal, Weibull, and Pareto families for the cost curves), and chose the least biased priors for each distribution from these. Across all Bayesian models, we used 4 Stan chains with 10000 burn-in iterations and 10000 sampling iterations. Bias and identifiability were examined through theoretic simulations using P-P plots [8]. We tested whether the chains achieved high coverage of the posterior distribution by checking effective sample size (N_eff_, where an N_eff_ of 10% of the iterations suggests unbiased sampling) and tested for chain convergence with the Gelman-Rubin diagnostic (R_hat_), using a threshold of 1.1, via the R package *shinystan* [9].

A note on prior choices

We tested uniform, Jeffreys, reference, and other simple prior formulations (e.g. inverse prior) and chose the least biased priors for each distribution from these. The Jeffreys prior is defined as: $\boldsymbol{\pi}\left( \boldsymbol{\theta} \right)\boldsymbol{=}\sqrt{\boldsymbol{Det(I}\left( \boldsymbol{\theta} \right)\boldsymbol{)}}$, where I($\boldsymbol{\theta}$) is the Fisher information matrix. Jeffreys [10] developed the prior as a parameterization-invariant alternative to the uniform prior. The reference prior [11] can produce similar properties and sometimes has better behavior. However, some of these formulations are very difficult to compute, and so simpler formulations such as the inverse were substituted when theoretic analyses showed they behaved well.

Mortality model

We fit a beta distribution to our host severity frequency distribution. The beta distribution is described by two free parameters, a and b:

$$\boldsymbol{proportional mortality \sim Beta(a,b)}$$

We chose priors for a and b based on theoretical simulations, where we drew 1000 random a and b parameters via Latin hypercube sampling (Table S9) and subsequently sampled from each of these beta distributions to produce 100 severity estimates to use as data to fit our Bayesian models. To assess bias, we examined the resulting P-P plots for each model [8]. P-P plots allow for the checking of bias and uncertainty in the posterior distribution by plotting the percentiles of the posterior distribution under which the true parameters lie. The percentiles should follow the 1:1 line in a purely unbiased model. Deviations from the line can indicate over- or underestimation, as well as over- or under-prediction of uncertainty. We found that the best-behaved P-P plots corresponded to a prior of $\frac{1}{\sqrt{ab}}$ for the model (see Yang & Berger, *unpubl. manuscript*), though they still resulted in slight overestimation (Fig. S8).

We performed posterior checks on our fitted Stan model through *shinystan* [9] to determine whether a tractable model could be estimated using uncertain, sequential bounds and no point mortality estimates. Our fitted model generated no warnings for the standard posterior checks (i.e., effective sample size N_eff_, Gelman-Rubin diagnostic R_hat_).

Mortality model results

The host mortality distribution groups pest-host combinations into a series of sequential bins based on severity (Fig. 3,S9; Table S10; [6]). The two lowest bins have uncertain upper bounds, and the highest bin has an uncertain lower bound. Our model thus included the relative frequencies of species within each bin as data, and we fit parameters for the beta distribution shape and scale, as well as for the bounds of the two lowest bins and the highest bin. We assumed that the mortality categories did not overlap and that mortality increased in severity, such that these bins could be assigned sequentially. The likelihood was the sum of the integrals under the associated probability density functions for different beta parameter sets and threshold values, making the log likelihood:

$$\boldsymbol{LL=}\boldsymbol{A}_{\boldsymbol{spp}}\boldsymbol{(log}\int_{\boldsymbol{i=0}}^{\boldsymbol{AT}} \boldsymbol{p}\left( \boldsymbol{i} \right|\boldsymbol{a,b))+}\boldsymbol{B}_{\boldsymbol{spp}}\boldsymbol{(log}\int_{\boldsymbol{i=AT}}^{\boldsymbol{BT}} \boldsymbol{p}\left( \boldsymbol{i} \right|\boldsymbol{a,b))+}\boldsymbol{C}_{\boldsymbol{spp}}\boldsymbol{(log}\int_{\boldsymbol{i=BT}}^{\boldsymbol{0.01}} \boldsymbol{p}\left( \boldsymbol{i} \right|\boldsymbol{a,b))+}\boldsymbol{D}_{\boldsymbol{spp}}\boldsymbol{(log}\int_{\boldsymbol{i=0.01}}^{\boldsymbol{0.1}} \boldsymbol{p}\left( \boldsymbol{i} \right|\boldsymbol{a,b))))+}\boldsymbol{E}_{\boldsymbol{spp}}\boldsymbol{(log}\int_{\boldsymbol{i=0.1}}^{\boldsymbol{0.25}} \boldsymbol{p}\left( \boldsymbol{i} \right|\boldsymbol{a,b))))+}\boldsymbol{F}_{\boldsymbol{spp}}\boldsymbol{(log}\int_{\boldsymbol{i=0.25}}^{\boldsymbol{0.95}} \boldsymbol{p}\left( \boldsymbol{i} \right|\boldsymbol{a,b))))+}\boldsymbol{G}_{\boldsymbol{spp}}\boldsymbol{(log}\int_{\boldsymbol{i=0.95}}^{\boldsymbol{GT}} \boldsymbol{p}\left( \boldsymbol{i} \right|\boldsymbol{a,b))))+}\boldsymbol{H}_{\boldsymbol{spp}}\boldsymbol{(log}\int_{\boldsymbol{i=GT}}^{\boldsymbol{1.0}} \boldsymbol{p}\left( \boldsymbol{i} \right|\boldsymbol{a,b))}$$

Where the letters *A-H* correspond to the binned severity categories listed in Fig. S9 and *i* is the proportional mortality. The posterior can then calculated via:

$$\boldsymbol{p}\left( \boldsymbol{a,b} | \boldsymbol{y} \right)\boldsymbol{\propto p}\left( \boldsymbol{a,b} \right)\boldsymbol{p(y|a,b)\propto-}\log\left( \boldsymbol{a} \right)\boldsymbol{-}\log\left( \boldsymbol{b} \right)\boldsymbol{+}\boldsymbol{A}_{\boldsymbol{spp}}\boldsymbol{(log}\int_{\boldsymbol{i=0}}^{\boldsymbol{AT}} \boldsymbol{p}\left( \boldsymbol{i} \right|\boldsymbol{a,b))+}\boldsymbol{B}_{\boldsymbol{spp}}\boldsymbol{(log}\int_{\boldsymbol{i=AT}}^{\boldsymbol{BT}} \boldsymbol{p}\left( \boldsymbol{i} \right|\boldsymbol{a,b))+}\boldsymbol{C}_{\boldsymbol{spp}}\boldsymbol{(log}\int_{\boldsymbol{i=BT}}^{\boldsymbol{0.01}} \boldsymbol{p}\left( \boldsymbol{i} \right|\boldsymbol{a,b))+}\boldsymbol{D}_{\boldsymbol{spp}}\boldsymbol{(log}\int_{\boldsymbol{i=0.01}}^{\boldsymbol{0.1}} \boldsymbol{p}\left( \boldsymbol{i} \right|\boldsymbol{a,b))))+}\boldsymbol{E}_{\boldsymbol{spp}}\boldsymbol{(log}\int_{\boldsymbol{i=0.1}}^{\boldsymbol{0.25}} \boldsymbol{p}\left( \boldsymbol{i} \right|\boldsymbol{a,b))))+}\boldsymbol{F}_{\boldsymbol{spp}}\boldsymbol{(log}\int_{\boldsymbol{i=0.25}}^{\boldsymbol{0.95}} \boldsymbol{p}\left( \boldsymbol{i} \right|\boldsymbol{a,b))))+}\boldsymbol{G}_{\boldsymbol{spp}}\boldsymbol{(log}\int_{\boldsymbol{i=0.95}}^{\boldsymbol{GT}} \boldsymbol{p}\left( \boldsymbol{i} \right|\boldsymbol{a,b))))+}\boldsymbol{H}_{\boldsymbol{spp}}\boldsymbol{(log}\int_{\boldsymbol{GT}}^{\boldsymbol{1.0}} \boldsymbol{p}\left( \boldsymbol{i} \right|\boldsymbol{a,b))}$$

Appendix S4. Additional sources of uncertainty and limitations

The major additional sources of uncertainty that were not examined in this modelling framework include 1) future climatic variability, 2) spread model uncertainty, 3) host distributional model uncertainty, 4) variability in management behavior, 5) asymptotic mortality misestimation, and 6) mortality debt model misspecification. In Table S9, we have categorized the relative level of uncertainty across land types associated with the different impact metrics reported in this paper. The table includes all of the elements from the above list impacting each of these categories. All impact metrics are sensitive to climatic variability, as well as a correctly-specified pest spread and tree distributional models. Our pest spread model is demonstrably predictive and consistent across land types, but we are more confident in our urban tree models for street trees than residential and community trees, as we were able to draw on a much larger fitting dataset for these, so all impact metrics have greater uncertainty for residential and community trees. Our most robust impact metric is predicted focal host exposure to each pest, since it is only sensitive to pest spread and urban tree distributional models. Asymptotic mortality refers to the maximum mortality predicted from our mortality model that can be attributed to forecasted pest spread from 2020 to 2050. This mortality would be reached eventually in any mortality debt scenario, and relies on accurate mortality estimates from our Bayesian approach. Specifying a time-window of 2020 to 2050 for impacts make the final two metrics (mortality and cost 2020 to 2050) more sensitive to the correct mortality debt model, but we know that the more uncertain pest species in terms of mortality debt scenario contribute less substantially to cost estimates. As such, we are relatively confident in all mortality estimates reported for 2020 to 2050 across all three land types. In addition to these sources of uncertainty, management responses cause additional uncertainty when translating from mortality within a given time range and cost within that range. These behaviors are most certain for street tree management, moderately uncertain for residential tree management, and most uncertain for community tree management (Table S14).

Non-street trees are likely at a lower priority for removal, and even within these other uses, management may differ. For example, dead trees may be left standing in wooded areas, but that is less likely in heavily used public parks where such trees would be hazardous to park users. Though we report rough cost estimates for the cases where these other trees are managed analogously to street trees (all dead trees are removed, taking into account that homeowners must pay more for tree treatment compared to municipalities who can benefit from economies of scale, [3]), we believe this is likely an overestimate, because responsibility and behavior is likely far more variable. Indeed, this may be a particularly severe overestimate for non-residential community trees, as management jurisdictions may be ambiguous for these trees, and many may not put people at risk (e.g., trees deep in wooded areas of parks or in industrial areas). While homeowners are likely to respond more frequently to dead trees, residential management is likely still to be more variable than street tree management. For instance, residential trees will likely be cut and replaced if the risk of them falling on a home is very high, but could be ignored otherwise, and this decision is likely to vary with variables such as lot size and household income. Nonetheless, the sheer number of projected deaths of non-street trees likely means they will contribute substantially to future costs.

We note that all known host genera for invasive forest pests were present in at least one community within our fitting set [7,12]. However, there are clear spatial biases in our records (Fig. S3). The majority of the communities in our fitting set are in the northern US and California. Our data spanned 32 states, but include relatively few communities in the southern US. It is possible that the regions we were missing have different relationships with the modelled predictor variables, but we were limited to the data available. Nonetheless, our data spanned a variety of climates (plant hardiness zones 3-11,[13]) and our communities ranged widely in population size (70 people – 8.6M people), which likely indicates a variety of potential tree assemblages.

We also note that the database upon which we built our mortality models [6] is based partly on expert elicitation, and is subject both to a high degree of imprecision and a potential bias toward overestimation of pest risk from forestry experts who have witnessed low-probability events [14]. Fei et al. [15] recently published empirical mortality rates for a subset of these pest species based on forested area data (Forest Inventory and Analysis program, (FIA), [12]), which were reported relative to background forested host mortality, and included estimates for some pathogens. Had we included pathogens in our analysis, we could have underestimated the mortality rate for deadly pathogens such as laurel wilt disease compared to these authors [15]. However, mortality rates in forested stands may not align well with urban host mortality risk for these pest species [16]. Nonetheless, the characterization of future pathogen-induced urban tree impacts is an important area of complementary analysis given the potential for high mortality rates. Background tree mortality was also not accounted for in our models, which may have killed some exposed hosts before the pests could cause impacts, especially in the longer time-horizon mortality debt models, since natural mortality probabilities are likely high on the scale of 100 years. Since the majority of projected damage is due to shorter-term dynamics, background mortality should offset only a small fraction of future pest-induced tree mortality in the next 30 years.

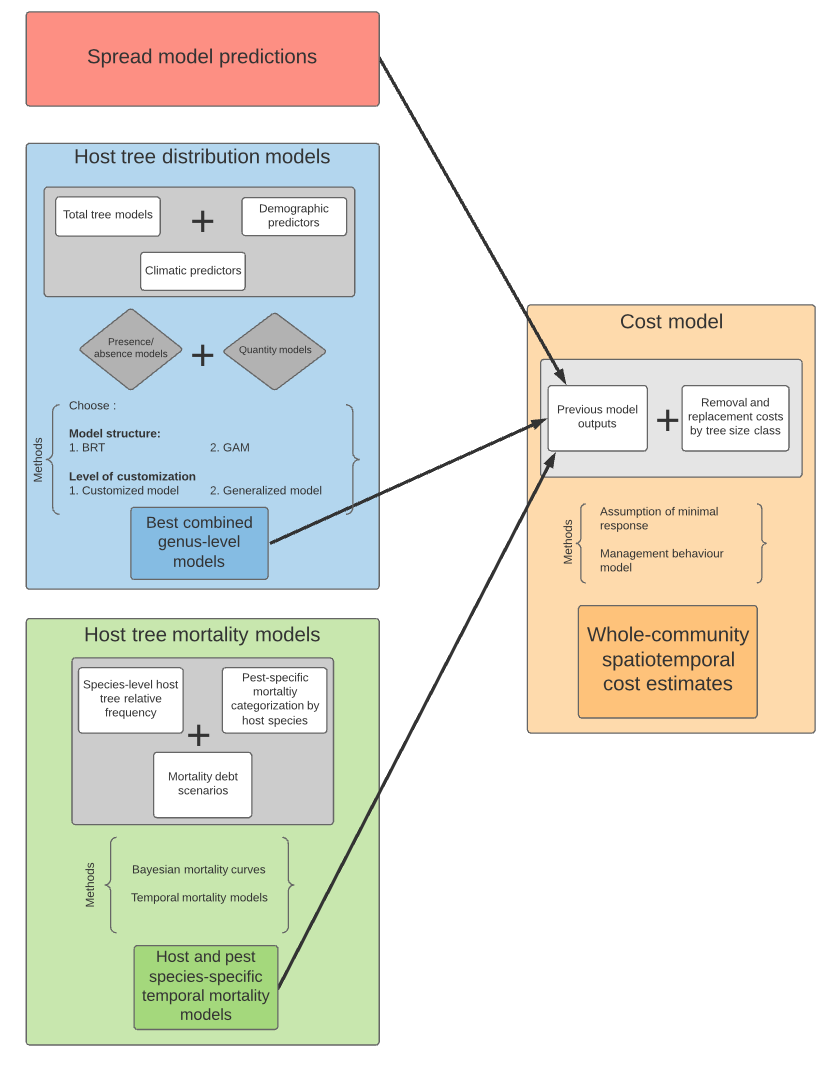

**Fig. S1.** Schematic representation of the four subcomponent models we combined to produce refined damage estimates to street trees from invasive US forest pests. Methodology is represented within braces, where GAM= Generalized Additive Model, and BRT= Boosted Regression Tree. The spread model predictions correspond to SDK forecasts from [17].

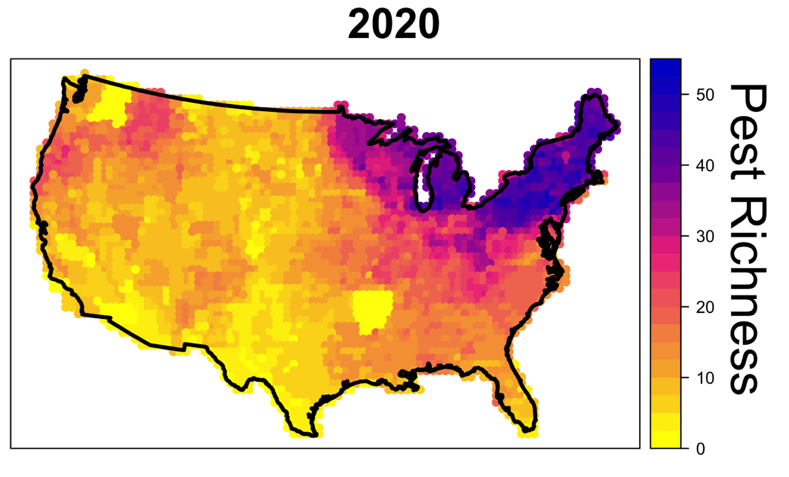

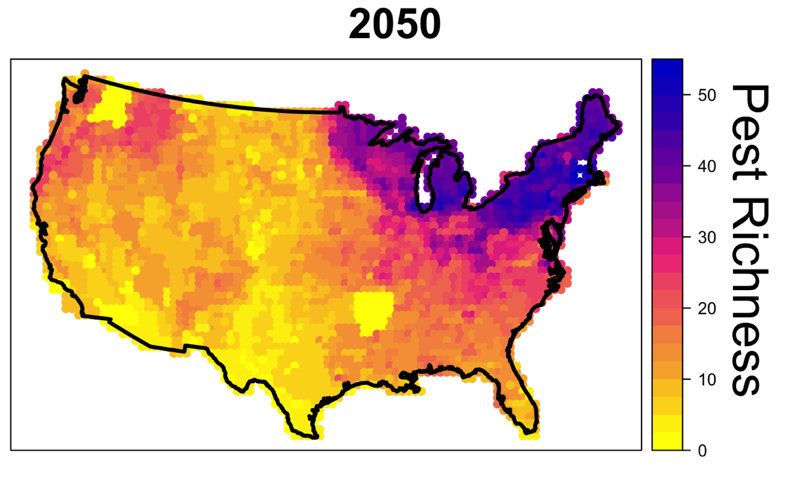

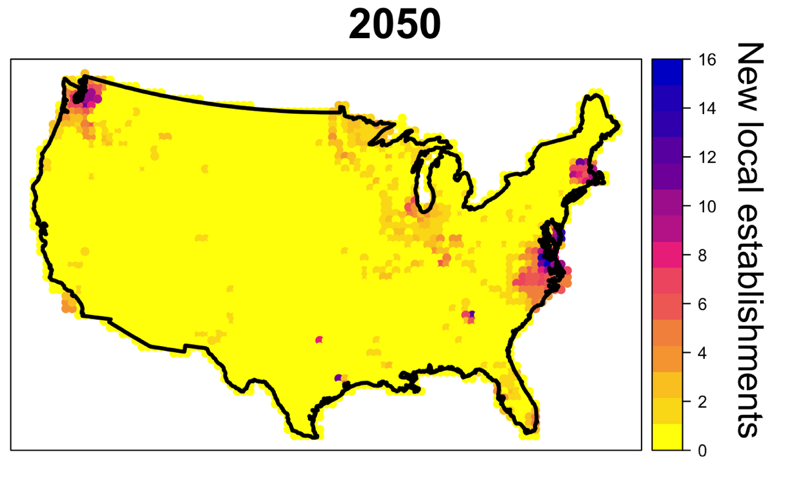

**Fig. S2.** Predicted pest richness in the mid-range climate scenario from 2020 to 2050, with newly occurring local establishments plotted in the last panel.

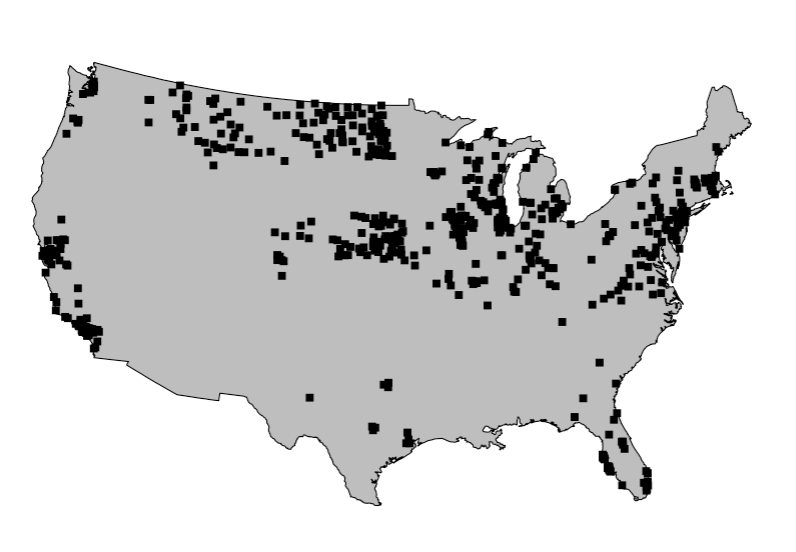

Fig. S3. Map of the 653 inventoried street tree communities across the United States.

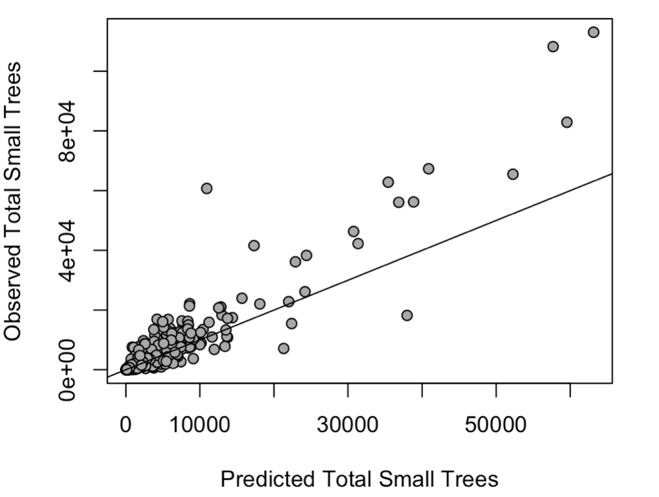

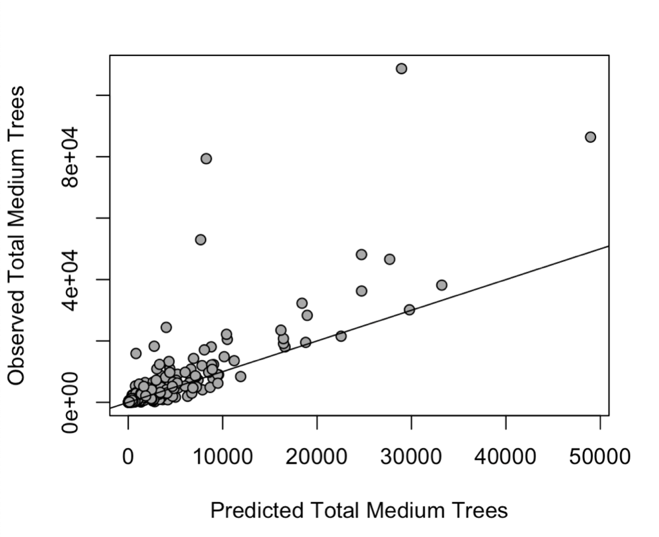

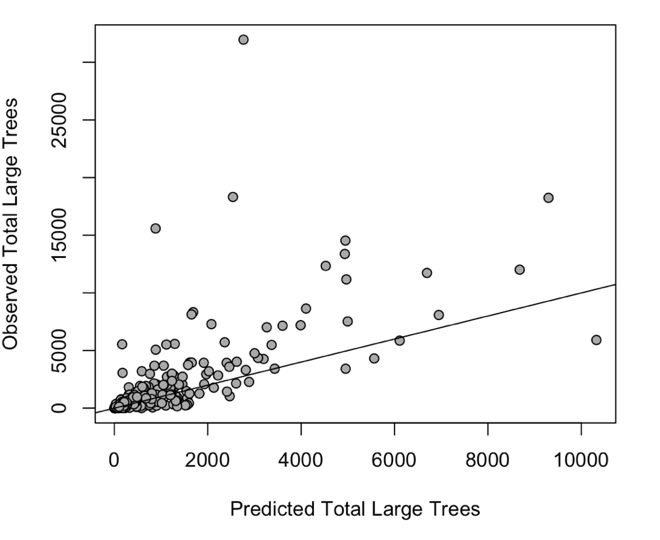

**Fig. S4.** Fits of the total tree abundance models to small, medium, and large trees across all 653 communities

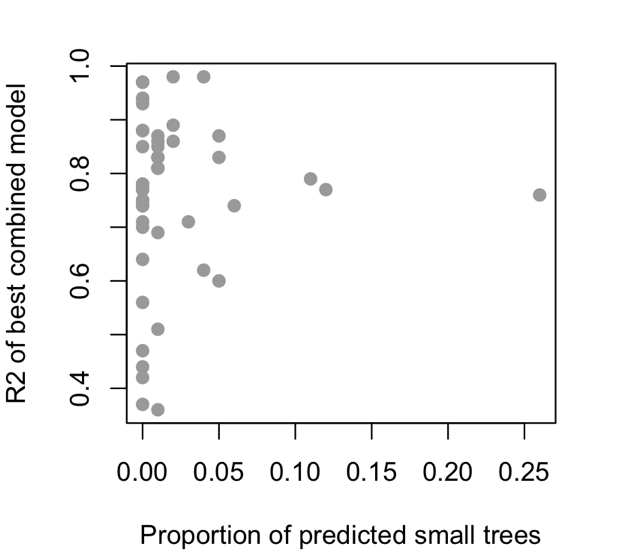

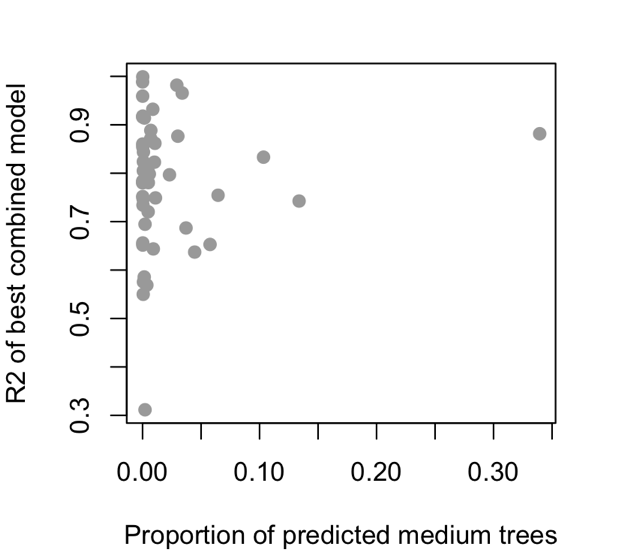

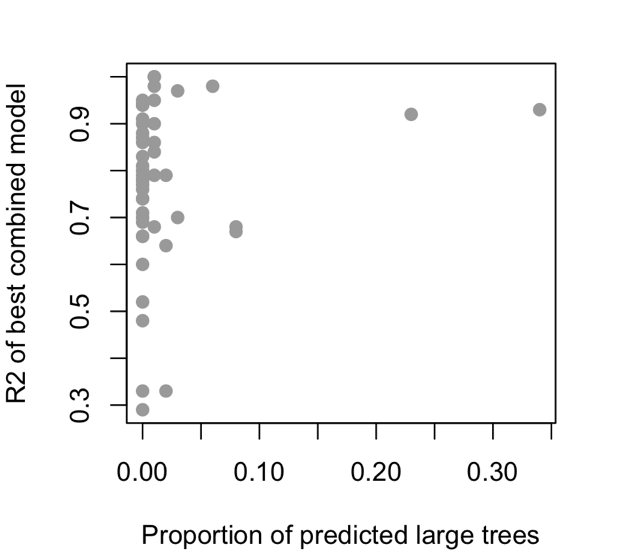

**Figure S5.** Inverse relationship between the rarity of a genus (defined as the predicted total number of trees of a particular size class) and the fit of the best single-genus model.

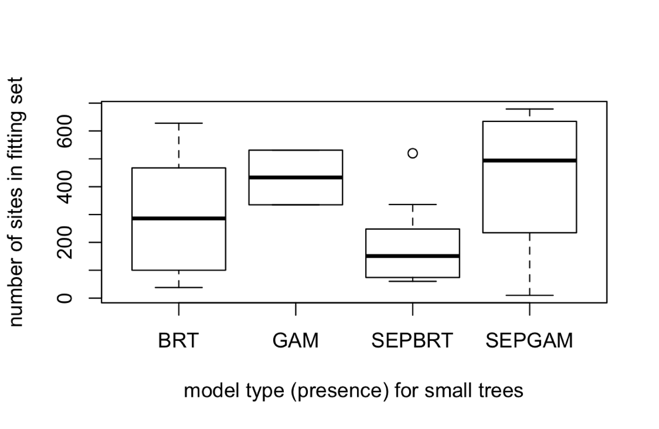

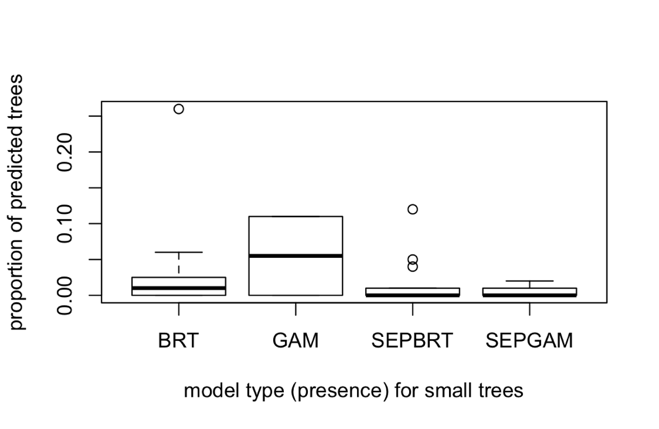

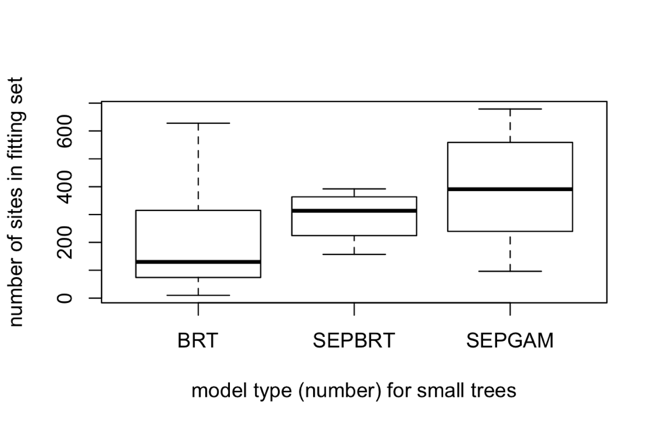

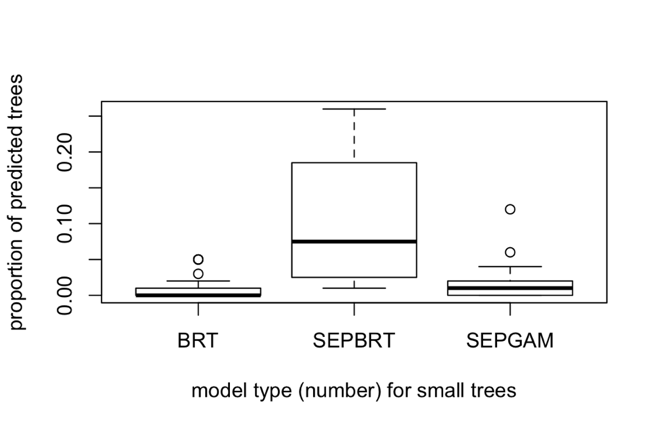

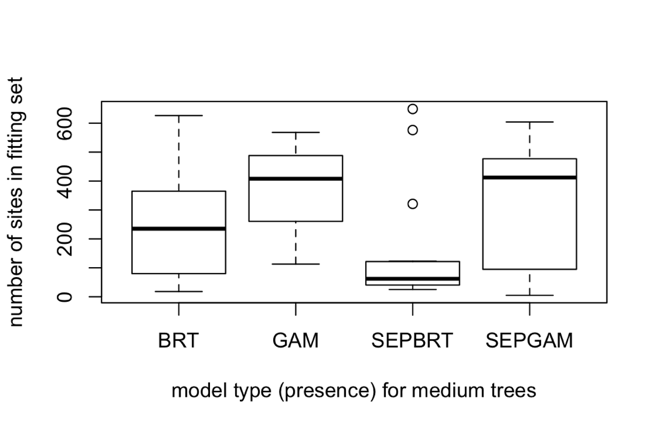

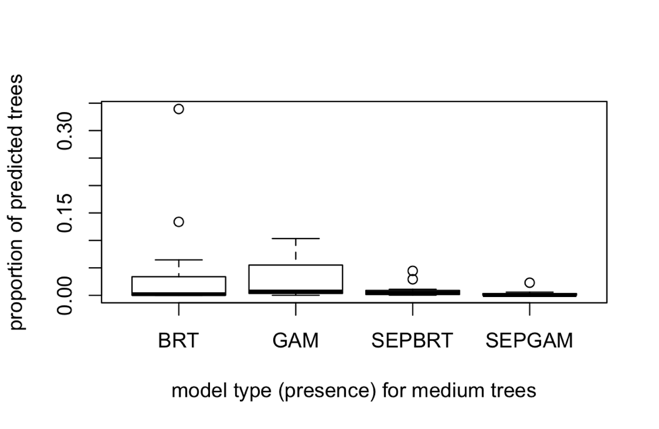

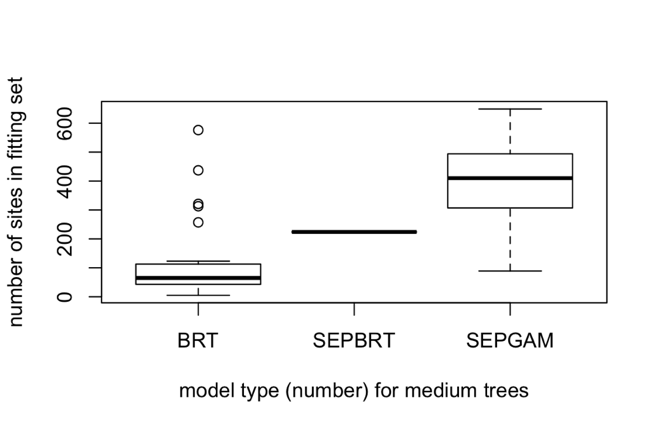

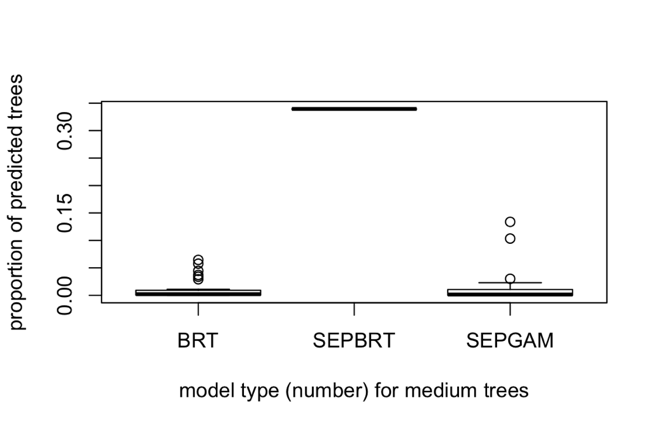

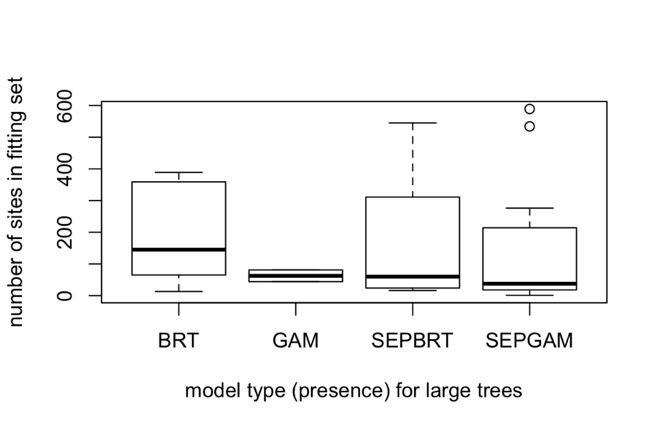

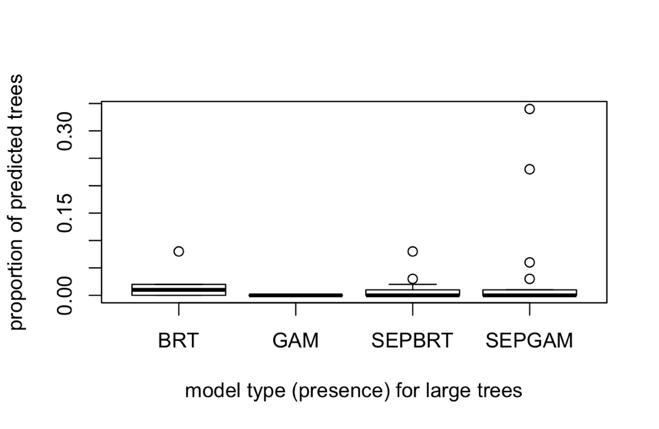

Model type (number) for small trees

Model type (presence) for small trees

Model type (presence) for medium trees

Number of sites in fitting set

Number of sites in fitting set

Model type (presence) for medium trees

Model type (number) for small trees

Model type (presence) for small trees

Proportion of predicted trees

Proportion of predicted trees

Proportion of predicted trees

Number of sites in fitting set

Number of sites in fitting set

Proportion of predicted trees

Model type (number) for medium trees

Model type (number) for medium trees

Proportion of predicted trees

Number of sites in fitting set

Model type (presence) for large trees

Model type (presence) for large trees

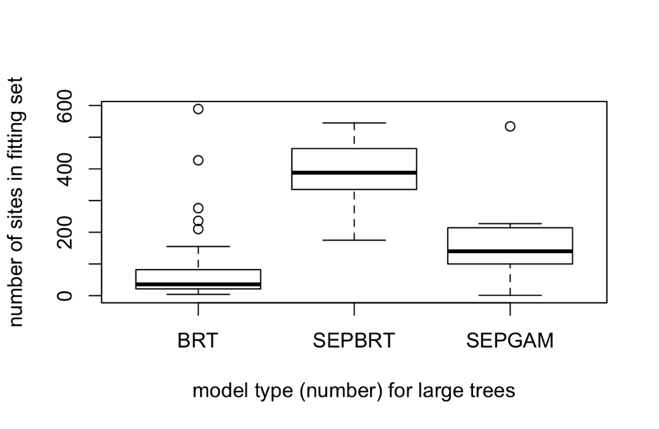

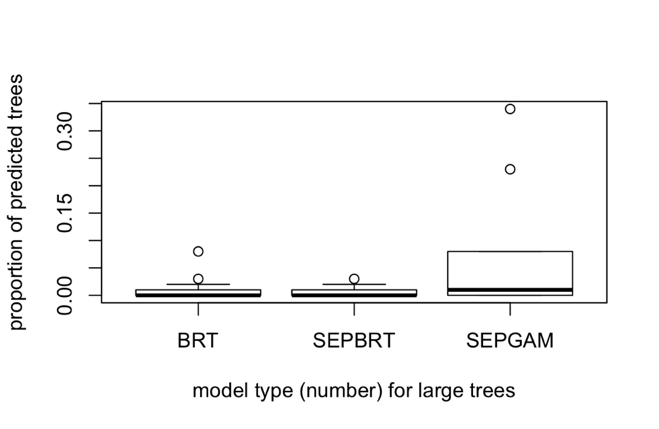

Model type (number) for large trees

Proportion of predicted trees

Number of sites in fitting set

Model type (number) for large trees

**Fig. S6.** Relationship between genus rarity (in terms of the number of sites where present and the predicted proportion of total trees) and the single-genus model selected across size classes. For small tree presence/absence, more GAMs and SEPGAMs are selected with more sites, while more GAMs are selected with more predicted trees. For small tree number, more SEPGAMs are selected with more sites, while more SEPBRTs are selected with more predicted trees. For medium tree presence/absence, fewer SEPBRTs are selected with fewer sites, while more BRTs and GAMs are selected with more predicted trees. For medium tree number, more SEPGAMs are selected for more sites, while more SEPBRTs are selected for more predicted trees. For large tree presence/absence, fewer GAMs are selected for fewer sites. For large tree number, more SEPBRTs are selected with more sites, and more SEPGAMs are selected with more predicted trees.

**a.	b.**

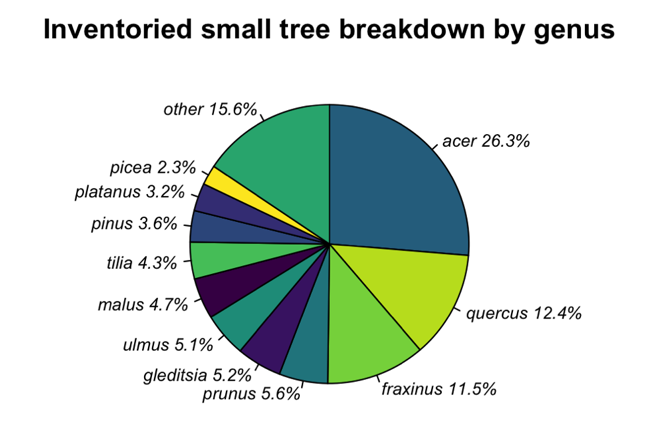

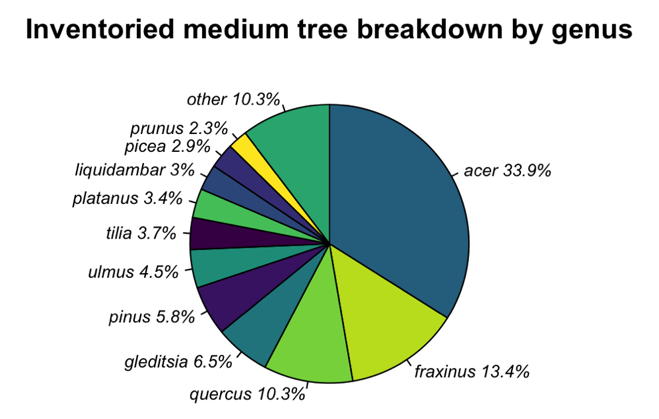

**c.	d.**

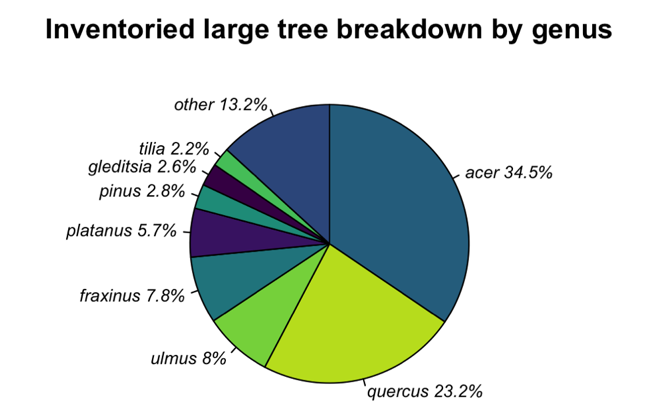

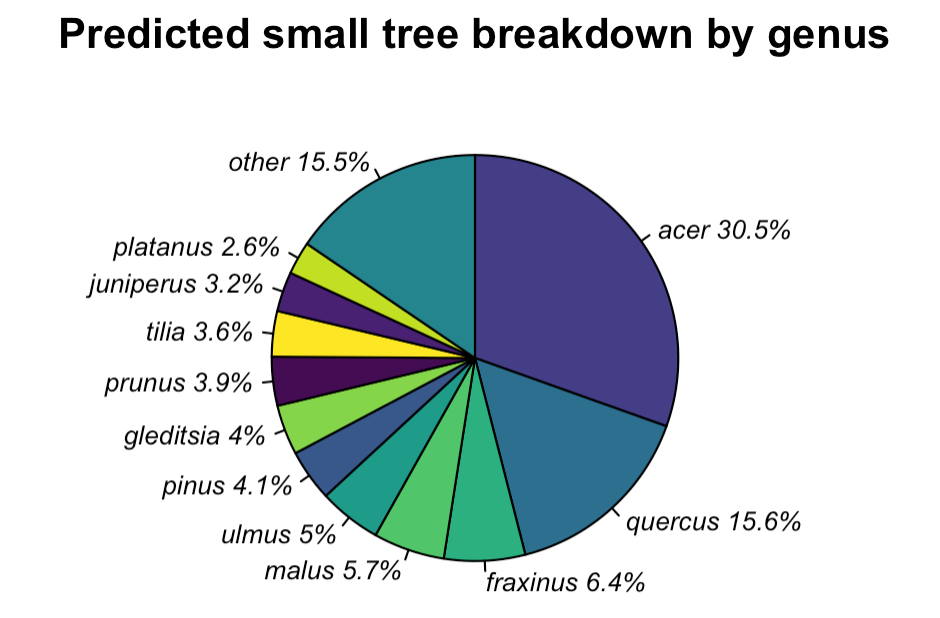
 **e.	f.**

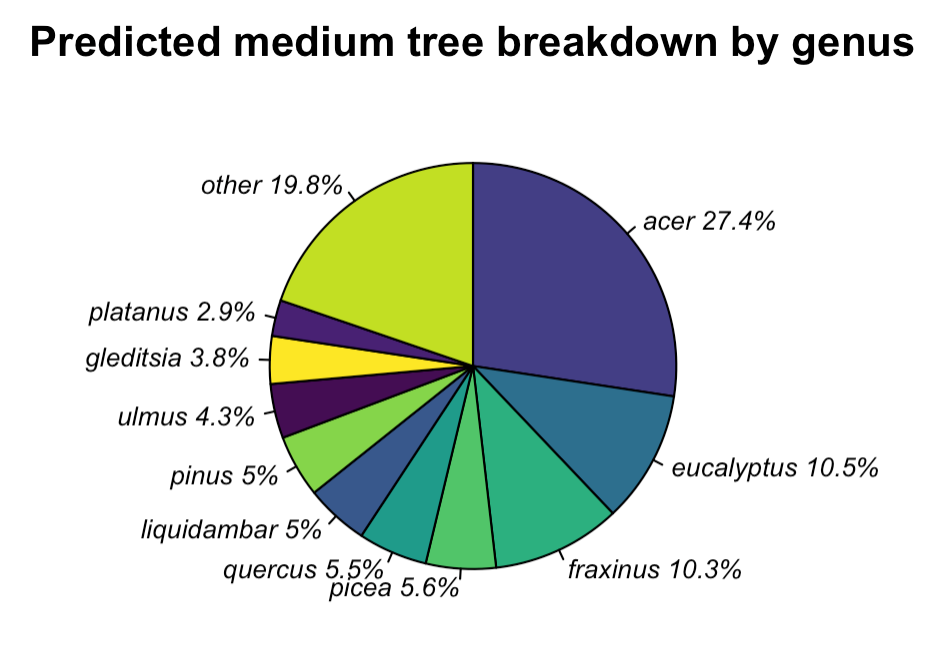

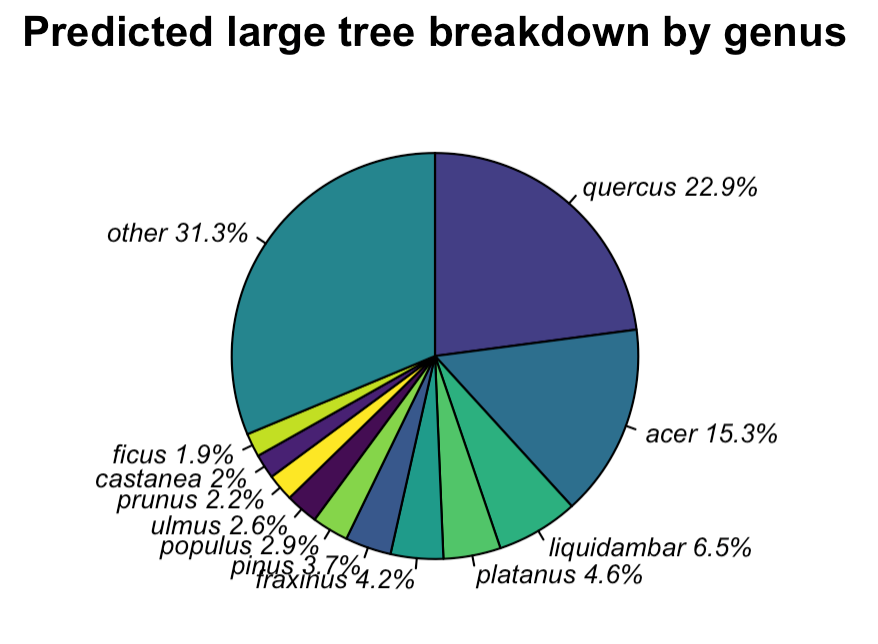

**Fig.S7.** Breakdown of both fitted (a-c) and extrapolated (d-f) small, medium and large tree abundance from the genus-specific models

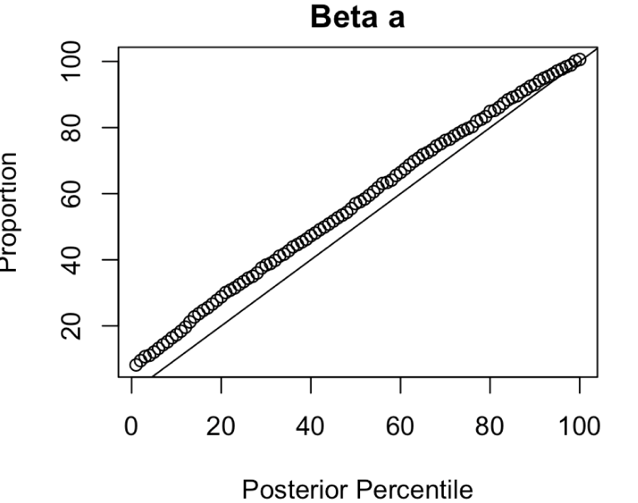

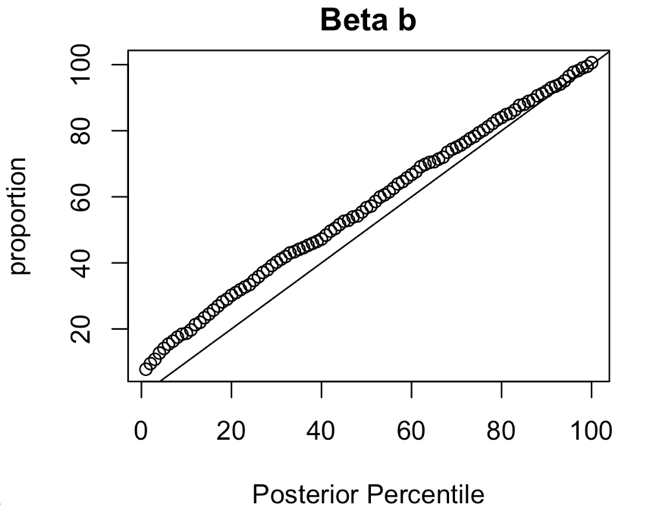

**Fig. S8.** PP-plots for the Latin hypercube sampling of our beta distribution model. Both parameters are slightly overestimated by the model, as evidenced by the positive deviation from the 1:1 line.

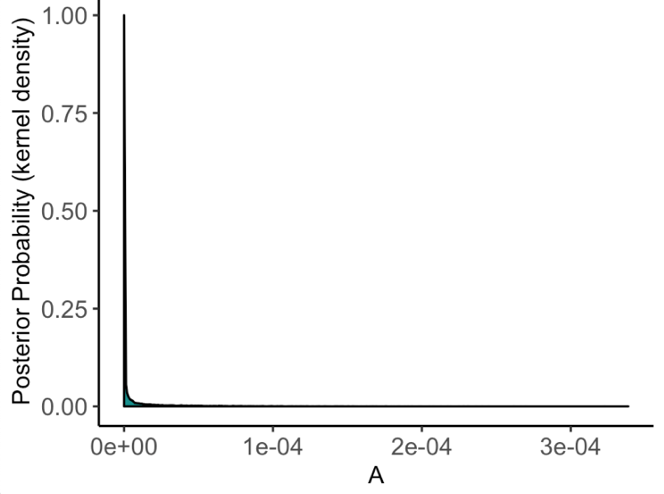

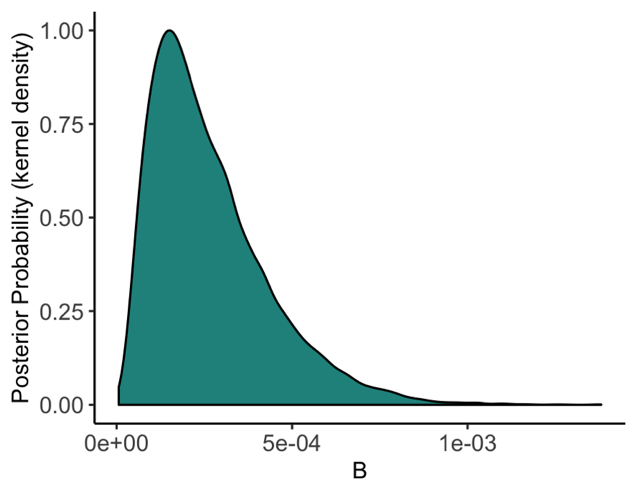

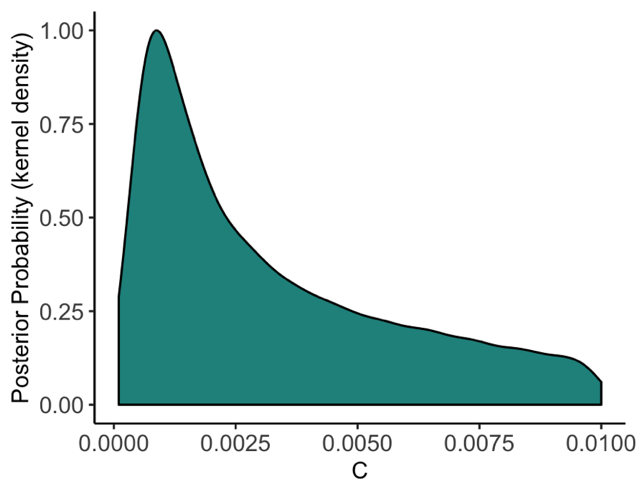

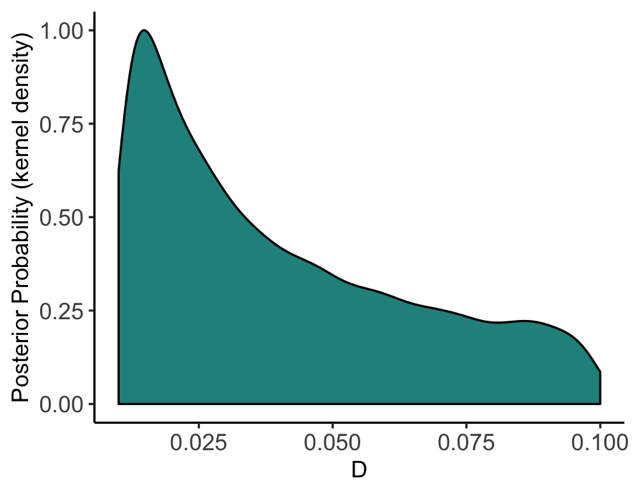

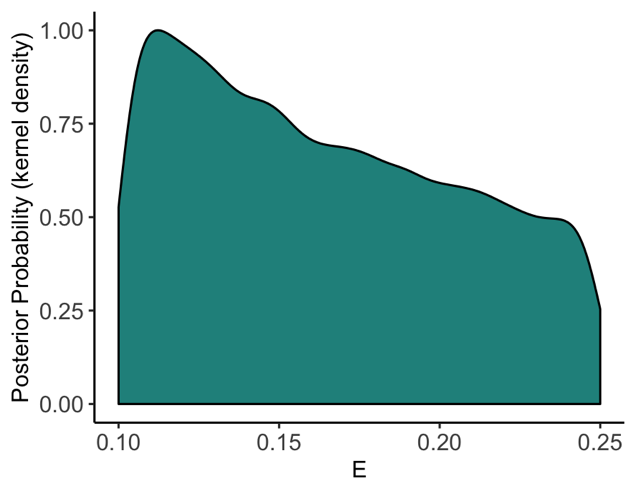

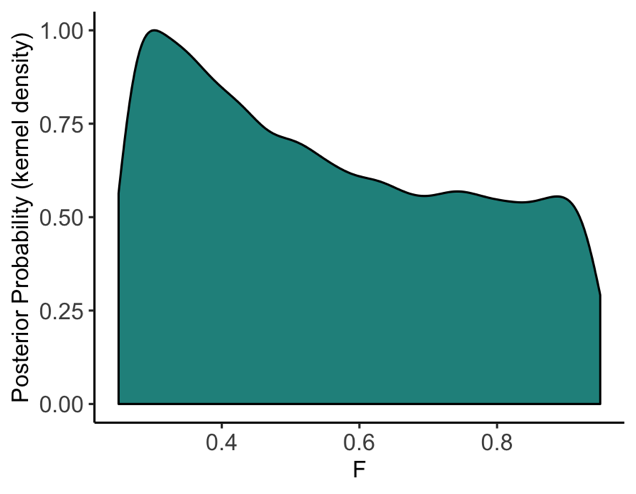

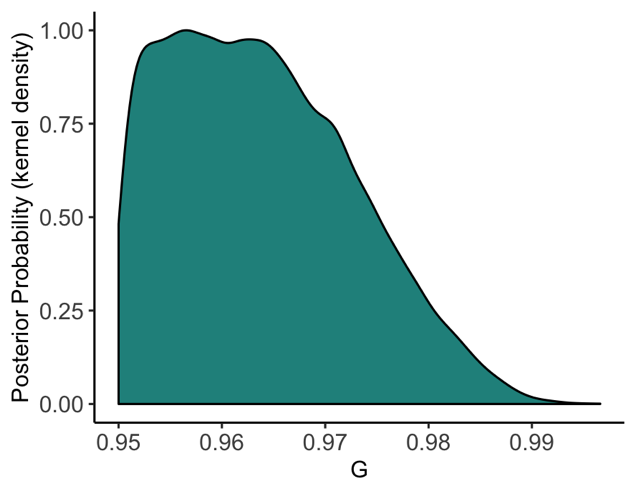

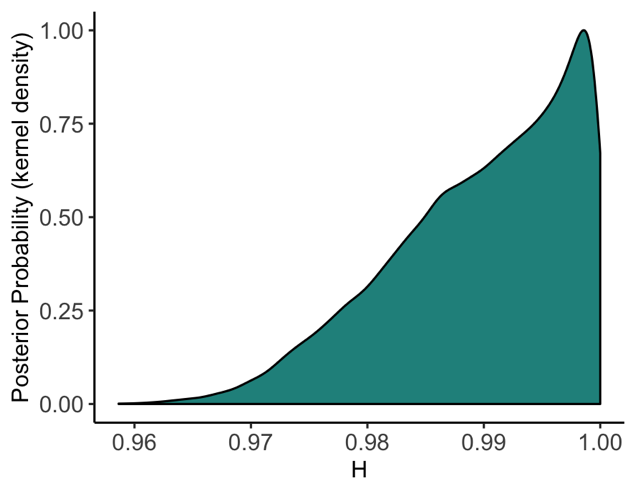

**Fig. S9.** Posterior distributions for the proportional mortality of pests in each severity category *A*=<<0.01, *B* = <0.01, *C*= 1-10%, *D*= 10-25%, *E*=25-95%, *F*=95-99%, *G*=99->99%, *H*= >99%-100%. These were sampled from to produce projections of host mortality due to each pest species across mortality debt and invasion lag scenarios.

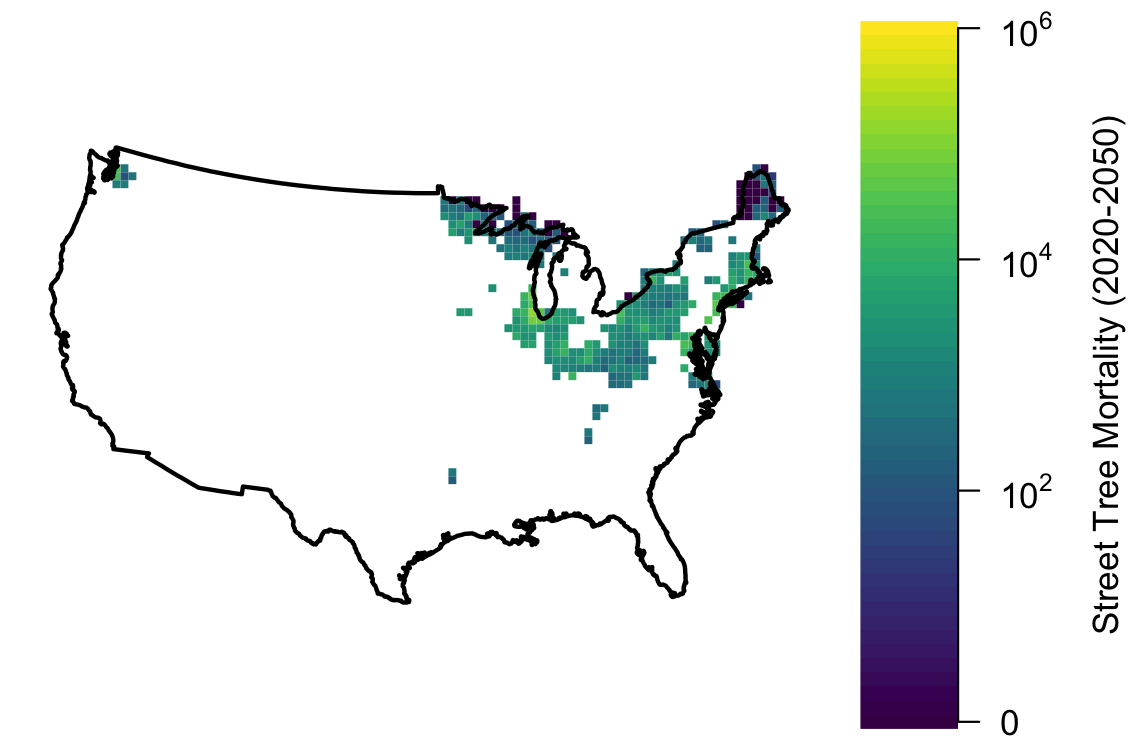

**Fig. S10.** Zone of asymptotic street tree mortality reached by EAB in the reasonable scenario..

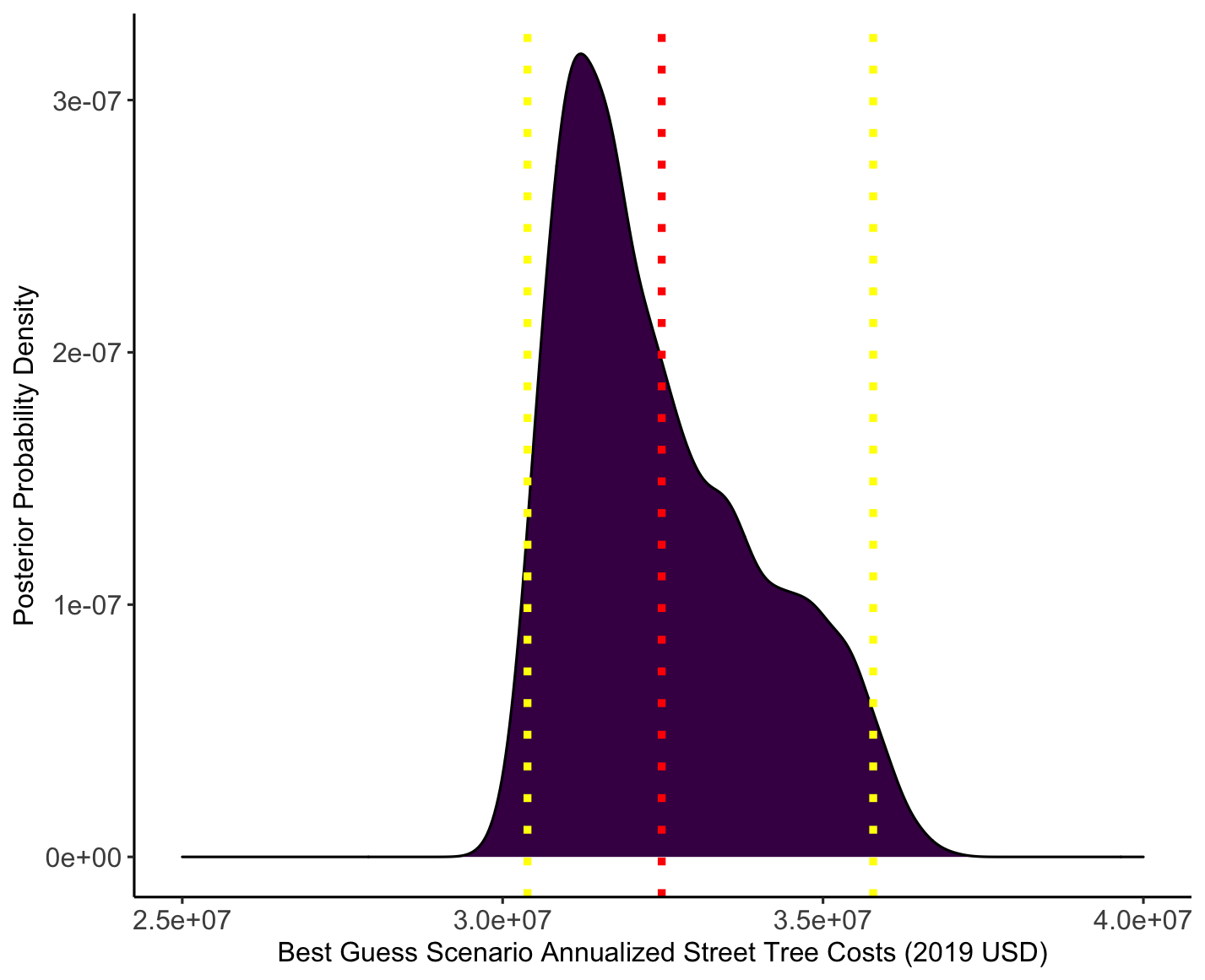
**Fig. S11.** Predicted annualized costs for the reasonable mortality debt scenario, with 95% Bayesian credible intervals shown in yellow and the posterior mean shown in red (46M USD).

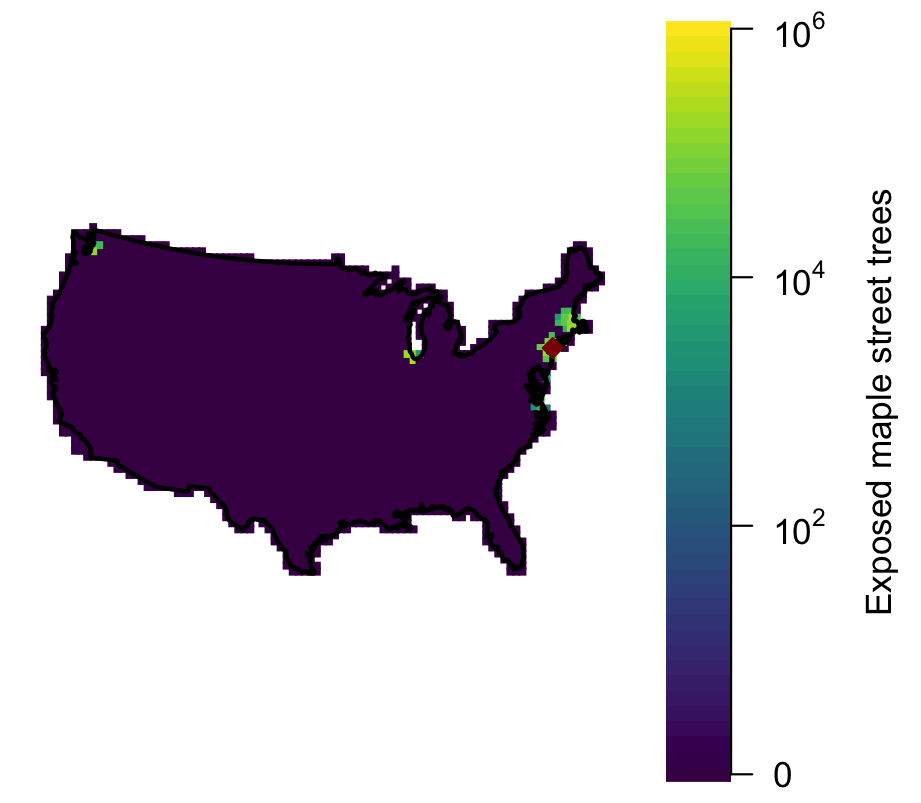

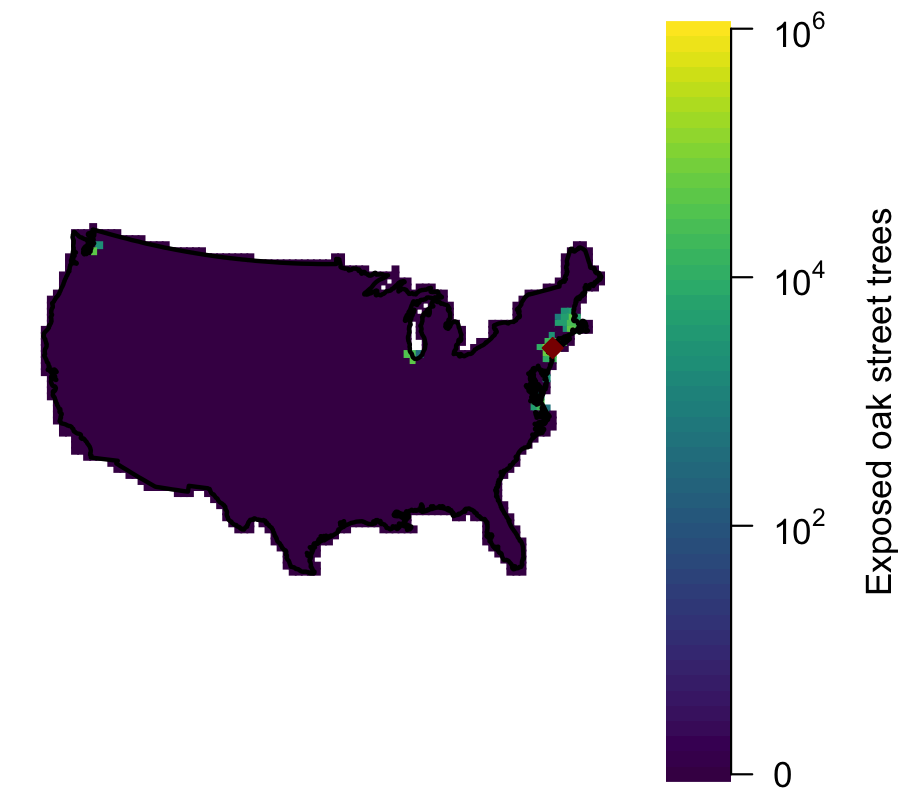

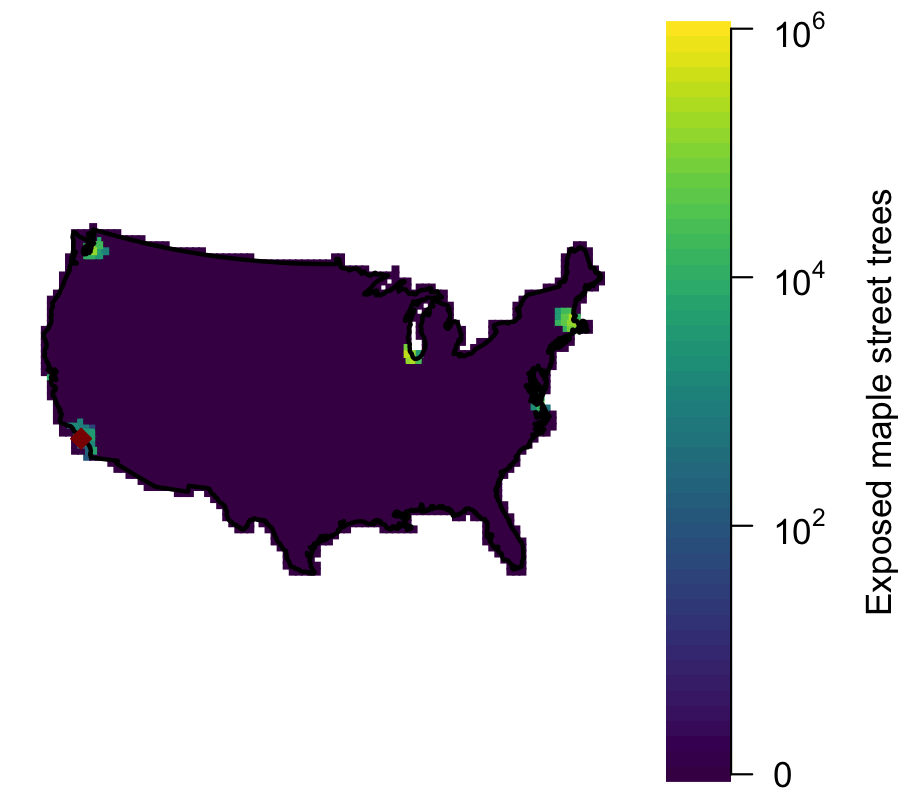

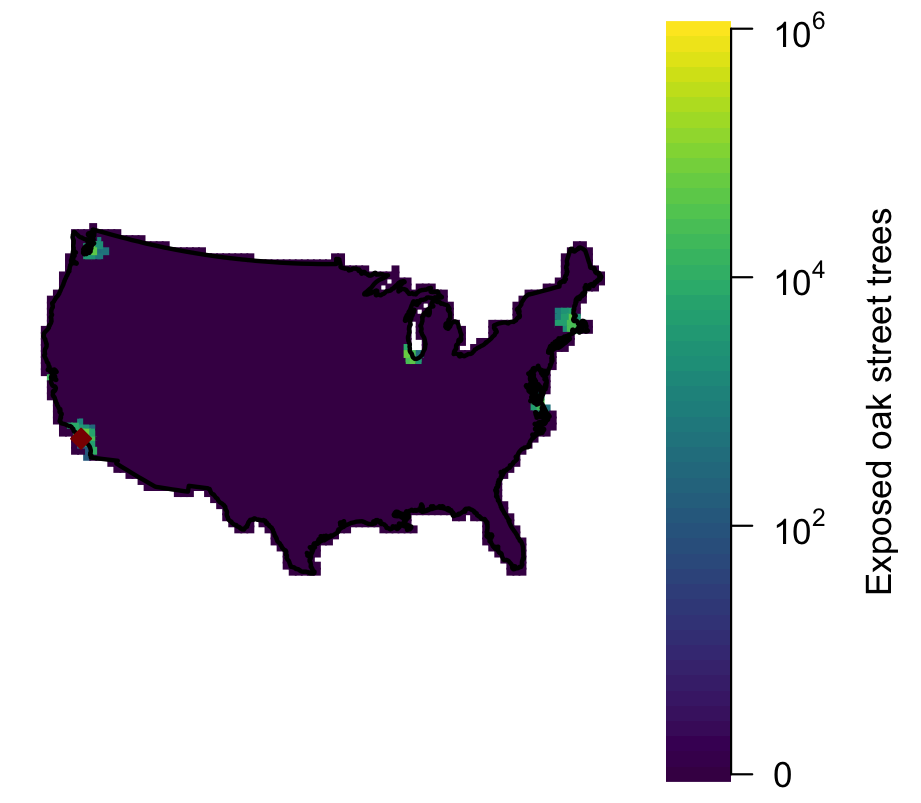

**f.**

**b.**

**d.**

**c.**

**a.**

**e.**

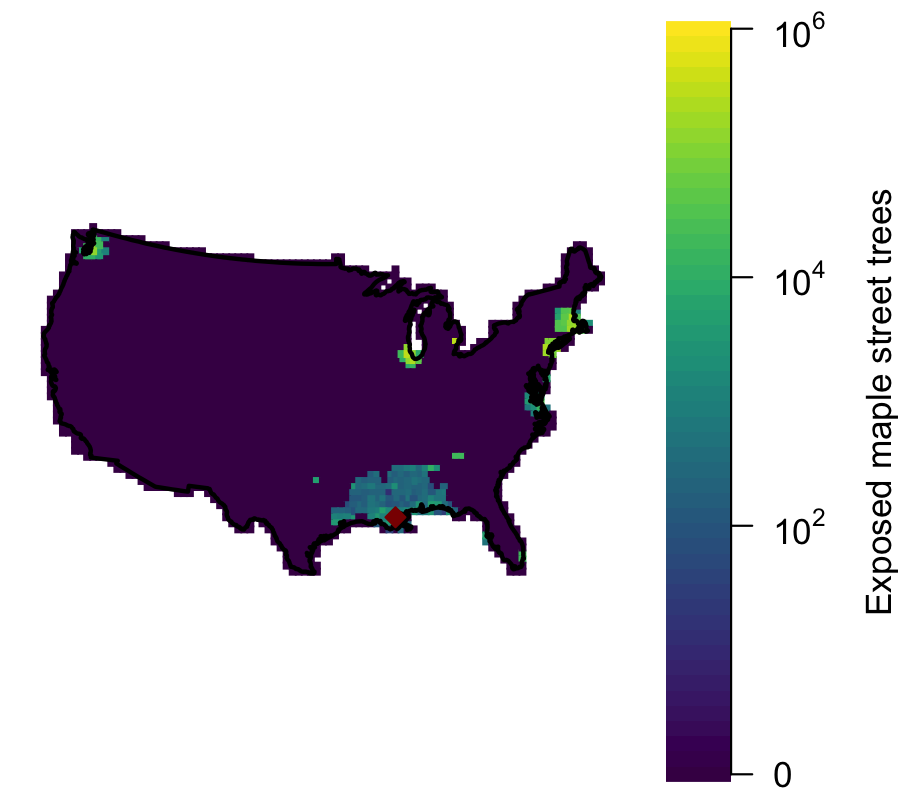

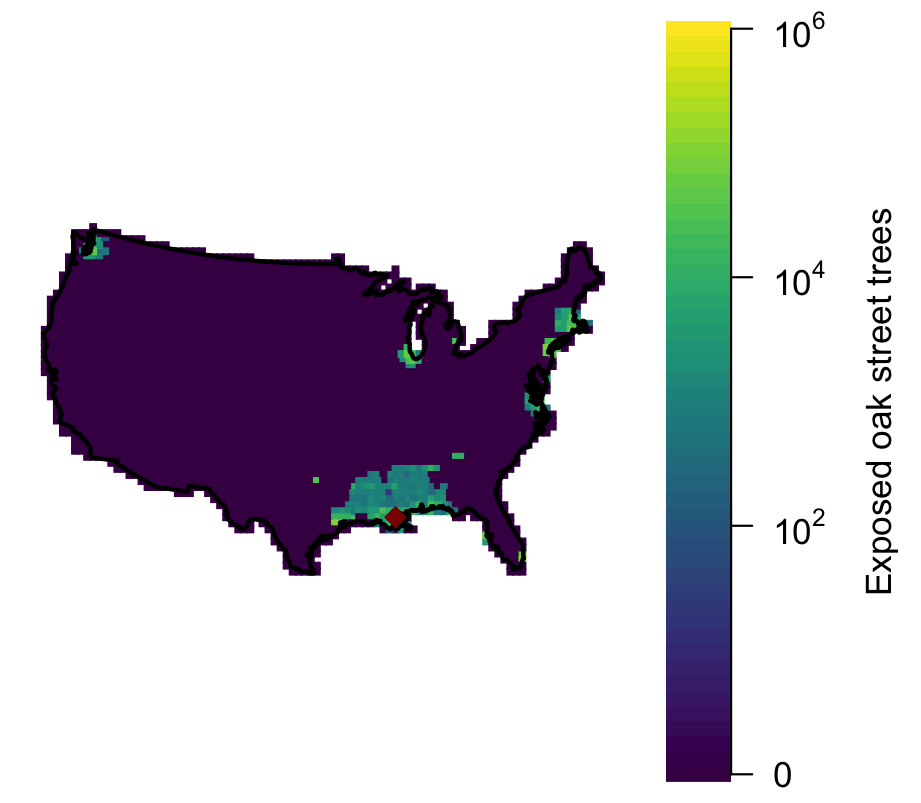

**Figure S12.** Resulting maple (**a,c,e**) and oak (**b,d,f**) exposure from 2020-2050 due to a simulated pest invasion originating the Port of New York and New Jersey (**a-b**), the Port of Long Beach (**c-d**)**,** and the Port of South Louisiana (**e-f**).

**Table S**1. List of IAFI species used in these analysis and their feeding guilds

| **Latin Name** | **Common Name** | **Feeding Guild** |
| --- | --- | --- |
| Agrilus planipennis | Emerald Ash Borer | Borers |
| Agrilus prionurus | Soapberry Borer | Borers |
| Anarsia lineatella | Peach Twig Borer | Borers |
| Anoplophora glabripennis | Asian Longhorned Beetle | Borers |
| Callidellum rufipenne | Japanese Cedar Longhorn Beetle | Borers |
| Cryptorhynchus lapathi | Poplar and Willow Borer | Borers |
| Hylastes opacus | European Bark Beetle | Borers |
| Hylurgus ligniperda | Red-haired Pine Bark Beetle | Borers |
| Orthotomicus erosus | Mediterranean Pine Engraver Beetle | Borers |
| Phoracantha recurva | Eucalyptus Longhorned Beetle | Borers |
| Scolytus multistriatus | Smaller European Elm Bark Beetle | Borers |
| Scolytus schevyrewi | Banded Elm Bark Beetle | Borers |
| Sirex noctilio | Sirex Wood Wasp | Borers |
| Tomicus piniperda | Pine Shoot Beetle | Borers |
| Acantholyda erythrocephala | Pine False Webworm | Defoliators |
| Coleophora laricella | Larch Casebearer | Defoliators |
| Contarinia baeri | European Pine Needle Midge | Defoliators |
| Cyrtepistomus castaneus | Asiatic Oak Weevil | Defoliators |
| Diprion similis | Introduced Pine Sawfly | Defoliators |
| Enarmonia formosana | Cherry Bark Tortrix | Defoliators |
| Epinotia nanana | European Spruce Needleminer | Defoliators |
| Euproctis chrysorrhoea | Browntail Moth | Defoliators |
| Fenusa pusilla | Birch Leafminer | Defoliators |
| Fenusa ulmi | Elm Leafminer | Defoliators |
| Gonipterus scutellatus | Eucalyptus Snout Beetle | Defoliators |
| Homadaula anisocentra | Mimosa Webworm | Defoliators |
| Leucoma salicis | Satin Moth | Defoliators |
| Lymantria dispar | LDD Moth | Defoliators |
| Neodiprion sertifer | European Pine Sawfly | Defoliators |
| Operophtera brumata | Winter Moth | Defoliators |
| Orchestes alni | Elm Flea Weevil | Defoliators |
| Otiorhynchus sulcatus | Black Vine Weevil | Defoliators |
| Plagiodera versicolora | Imported Willow Leaf Beetle | Defoliators |
| Popillia japonica | Japanese Beetle | Defoliators |
| Pristiphora erichsonii | Larch Sawfly | Defoliators |
| Pristiphora geniculata | Mountain Ash Sawfly | Defoliators |
| Profenusa thomsoni | Ambermarked Birch Leafminer | Defoliators |
| Rhyacionia buoliana | European Pine Shoot Moth | Defoliators |
| Trichiocampus viminalis | Poplar sawfly | Defoliators |
| Xanthogaleruca luteola | Elm Leafbeetle | Defoliators |
| Adelges abietis | Eastern Spruce Gall Adelgid | Sap-feeders |
| Adelges piceae | Balsam Woolly Adelgid | Sap-feeders |
| Adelges tsugae | Hemlock Woolly Adelgid | Sap-feeders |
| Aonidiella aurantii | California Red Scale | Sap-feeders |
| Asterolecanium variolosum | Golden Oak Scale | Sap-feeders |
| Blastopsylla occidentalis | Eucalyptus Psyllid | Sap-feeders |
| Carulaspis juniperi | Juniper Scale | Sap-feeders |
| Cryptoccocus fagisuga Lind. | Beech Scale | Sap-feeders |
| Diaspidiotus perniciosus | San Jose Scale | Sap-feeders |
| Elatobium abietinum | Green Spruce Aphid | Sap-feeders |
| Eulecanium cerasorum | Calico Scale | Sap-feeders |
| Fiorinia externa | Elongate Hemlock Scale | Sap-feeders |
| Glycaspis brimblecombei | Redgum Lerp Psyllid | Sap-feeders |
| Icerya purchasi | Cottony Cushion Scale | Sap-feeders |
| Lepidosaphes ulmi | Oystershell Scale | Sap-feeders |
| Maconellicoccus hirsutus | Pink Hibiscus Mealybug | Sap-feeders |
| Matsucoccus matsumurae | Red Pine Scale | Sap-feeders |
| Nuculaspis tsugae | Circular Hemlock Scale | Sap-feeders |
| Taeniothrips inconsequens | Pear Thrips | Sap-feeders |
| Thrips calcaratus | Introduced Basswood Thrips | Sap-feeders |

**Table S2.** List of predictors used in total tree and genus-specific tree models

| **Predictor** | **Unit** | **Description** | **Reference** |
| --- | --- | --- | --- |
| Population |  | 2010 Census population | US Census Bureau <http://www.census.gov/popest> |
| Ecological Province |  | Subregions using criteria defined in the National Hierarchical Framework of Ecological Units | [18] |
| Median number of freeze free days |  | Median number of days between last spring and first autumn temperature ≤0 °C | PRISM Climate Group, Oregon State University |
| Distance to Road | Log (km) | Mean distance to nearest road | US Geological Survey, Fort Collins Science Center; [5] |
| Total Road Length | Log (km^-1^) | Length of road per square km of community area | US Geological Survey, Fort Collins Science Center; [5] |
| Distance to Road * Total Road Length |  | See above | See above |
| Mean moisture index |  | Balance between precipitation and potential evapotranspiration; scaled between -1 and 1 | PRISM Climate Group, Oregon State University, [5,19] |
| Mean precipitation | mm | Mean annual measured precipitation | PRISM Climate Group, Oregon State University, [5,19] |
| Elevation | meters | Elevation above sea level | US Geological Survey National Elevation Dataset |
| Area | hectares | Community size | American Community Survey <https://www.census.gov/programs-surveys/acs> |
| Income | USD | Median household income in 1999 (County level) | US Census Bureau <http://quickfacts.census.gov/qfd/meta/long_INC910199.htm> |
| Mean Year of Home Construction | Year | Average age a home was built based on surveys from 2015  (block-group level) | American Community Survey  <https://www.census.gov/programs-surveys/acs> |
| Median Value of Home | USD | Median value of a home in the community in 2015  (block-group level) | American Community Survey  <https://www.census.gov/programs-surveys/acs> |
| bio2 | °C *10 | Mean Diurnal Range (Mean of monthly (max temp - min temp)) | WORLDCLIM [20] |
| bio8 | °C *10 | Mean Temperature of Wettest Quarter | WORLDCLIM [20] |
| bio10 | °C *10 | Mean Temperature of Warmest Quarter | WORLDCLIM [20] |
| bio11 | °C *10 | Mean Temperature of Coldest Quarter | WORLDCLIM [20] |
| bio13 | mm | Precipitation of Wettest Month | WORLDCLIM [20] |
| bio15 | mm^-2^ | Precipitation Seasonality (Coefficient of Variation) | WORLDCLIM [20] |
| Distance to coast | meters |  | Calculated in ArcGIS with US Equidistant Conic Projection |
| Tree canopy cover | % | Fraction of community area covered by trees | NLCD 2011 [21] |
| Latitude | meters |  | Calculated in ArcGIS with US Equidistant Conic Projection |
| Longitude | meters |  | Calculated in ArcGIS with US Equidistant Conic Projection |

**Table S3.** Results of total tree abundance models (a. small, b. medium, c. large). Models were fit via boosted regression trees, so relationships with predictors are not strictly positive or negative, but the overall shape of the relationship is summarized in the ‘general sign’ column.

| **Predictor** | **Relative Influence** | **General**  **Sign** |
| --- | --- | --- |
| **a. Small Trees** |  |  |
| Population | 55 | + |
| Area | 4.5 | + |
| Ecological province | 10.9 | NA |
| bio10 | 2.8 | - |
| Mean home value | 2.7 | + |
| Income | 2.2 | + |
| Mean year of home construction | 1.5 | - |
| Mean precipitation | 1.2 | + |
| Distance to road* Total road length | 1.2 | - |
| Others | <1 |  |
| **b. Medium Trees** |  |  |
| Population | 58.6 | + |
| Ecological province | 13.5 | NA |
| Area | 6.3 | + |
| Mean year of home construction | 5.8 | - |
| bio10 | 2.0 | parabolic (+, then – after inflection) |
| Income | 1.8 | + |
| Distance to road | 1.6 | - |
| Mean number of freeze free days | 1.4 | + |
| Total road length | 1.3 | - |
| bio11 | 1.2 | - |
| Mean home value | 1.2 | + |
| Distance to road * Total road length | 1.0 | - |
| Others | <1 |  |
| **c. Large trees** |  |  |
| Population | 43 | + |
| Mean year of home construction | 13.8 | - |
| Ecological province | 13 | NA |
| Area | 6.6 | + |
| Tree canopy cover | 6.5 | + |
| Mean number of freeze free days | 3.8 | + |
| bio11 | 3.4 | + |
| Distance to road | 1.2 | - |
| Mean precipitation | 1.1 | parabolic (positive then negative) |
| Income | 1 | + |
| Others | <1 |  |

**Table S4.** Strength of predictive ability across all single-genus tree models, with the selected best-fitting tree presence/absence and tree number model components shown for each (SEP indicates a model fit with genus-specific terms).

| **Genus** | **R^2^ (small)** | **Presence model (small)** | **Abundance model (small)** | **R^2^ (medium)** | **Presence model (medium)** | **Abundance model (medium)** | **R^2^ (large)** | **Abundance model (large)** | **Abundance model (large)** |
| --- | --- | --- | --- | --- | --- | --- | --- | --- | --- |
| *Abies* | 0.70 | BRT | BRT | 0.78 | SEPBRT | BRT | 0.68 | SEPGAM | BRT |
| *Acacia* | 0.37 | BRT | BRT | 0.78 | BRT | BRT | 0.94 | SEPGAM | BRT |
| *Acer* | 0.76 | BRT | BRT | 0.81 | SEPBRT | SEPGAM | 0.71 | SEPGAM | BRT |
| *Aesculus* | 0.74 | BRT | BRT | 0.82 | BRT | SEPGAM | 0.98 | BRT | BRT |
| *Amelanchier* | 0.87 | BRT | BRT | 0.31 | SEPBRT | BRT | 0.29 | SEPBRT | BRT |
| *Arbutus* | 0.85 | BRT | BRT | 0.81 | SEPBRT | BRT | 0.999 | SEPGAM | BRT |
| *Betula* | 0.86 | SEPGAM | SEPGAM | 0.86 | BRT | SEPGAM | 0.90 | BRT | SEPGAM |
| *Castanea* | 0.77 | BRT | BRT | 0.75 | BRT | BRT | 0.97 | SEPGAM | BRT |
| *Chamaecyparis* | 0.77 | BRT | BRT | 0.98 | SEPBRT | BRT | 0.90 | SEPBRT | BRT |
| *Cinnamomum* | 0.83 | BRT | BRT | 0.75 | SEPBRT | BRT | 0.87 | GAM | BRT |
| *Citrus* | 0.88 | BRT | BRT | 0.59 | SEPBRT | BRT | 0.94 | SEPGAM | BRT |
| *Cornus* | 0.81 | BRT | BRT | 0.99 | BRT | BRT | 0.48 | SEPBRT | BRT |
| *Cotinus* | 0.71 | SEPGAM | SEPGAM | 0.57 | BRT | BRT | 0.999 | SEPGAM | SEPGAM |
| *Crataegus* | 0.83 | BRT | BRT | 0.92 | GAM | BRT | 0.91 | SEPBRT | BRT |
| *Cupressus* | 0.75 | BRT | BRT | 0.69 | BRT | BRT | 0.79 | SEPBRT | BRT |
| *Elaeagnus* | 0.78 | BRT | BRT | 0.96 | BRT | SEPGAM | 0.77 | SEPGAM | BRT |
| *Eucalyptus* | 0.64 | BRT | BRT | 0.85 | BRT | SEPGAM | 0.70 | GAM | BRT |
| *Fagus* | 0.78 | BRT | BRT | 0.65 | BRT | SEPGAM | 0.74 | SEPBRT | BRT |
| *Ficus* | 0.97 | BRT | BRT | 0.78 | SEPGAM | BRT | 0.88 | BRT | BRT |
| *Fraxinus* | 0.79 | SEPBRT | SEPBRT | 0.82 | BRT | SEPGAM | 0.78 | SEPBRT | SEPBRT |
| *Gleditsia* | 0.83 | SEPBRT | SEPBRT | 0.86 | BRT | SEPGAM | 0.83 | SEPBRT | SEPBRT |
| *Ilex* | 0.81 | BRT | BRT | 0.551 | BRT | BRT | 0.52 | SEPBRT | BRT |
| *Juglans* | 0.44 | SEPGAM | SEPGAM | 0.91 | SEPBRT | BRT | 0.80 | SEPGAM | SEPGAM |
| *Juniperus* | 0.85 | SEPGAM | SEPGAM | 0.83 | GAM | SEPGAM | 0.92 | SEPGAM | SEPGAM |
| *Larix* | 0.74 | BRT | BRT | 0.64 | SEPBRT | BRT | 0.67 | SEPBRT | BRT |
| *Liquidambar* | 0.89 | SEPGAM | SEPGAM | 0.74 | BRT | SEPGAM | 0.68 | BRT | SEPGAM |
| *Liriodendron* | 0.51 | SEPBRT | SEPBRT | 0.65 | BRT | BRT | 0.70 | SEPBRT | SEPBRT |
| *Maclura* | 0.47 | BRT | BRT | 0.72 | SEPBRT | BRT | 0.79 | SEPGAM | BRT |
| *Magnolia* | 0.86 | SEPGAM | SEPGAM | 0.88 | BRT | SEPBRT | 0.93 | SEPGAM | SEPGAM |
| *Malus* | 0.87 | BRT | BRT | 0.80 | SEPGAM | SEPGAM | 0.84 | BRT | BRT |
| *Morus* | 0.69 | BRT | BRT | 0.75 | BRT | SEPGAM | 0.76 | SEPGAM | BRT |
| *Ostrya* | 0.42 | BRT | BRT | 0.64 | SEPBRT | BRT | 0.33 | BRT | BRT |
| *Persea* | 0.93 | BRT | BRT | 0.81 | SEPBRT | BRT | 0.74 | SEPGAM | BRT |
| *Picea* | 0.98 | BRT | BRT | 0.92 | SEPGAM | SEPGAM | 0.86 | SEPGAM | BRT |
| *Pinus* | 0.62 | SEPBRT | SEPBRT | 0.87 | GAM | SEPGAM | 0.66 | BRT | SEPBRT |
| *Platanus* | 0.71 | SEPBRT | SEPBRT | 0.88 | BRT | SEPGAM | 0.64 | BRT | SEPBRT |
| *Populus* | 0.36 | BRT | BRT | 0.80 | SEPGAM | SEPGAM | 0.95 | SEPBRT | BRT |
| *Prunus* | 0.74 | SEPGAM | SEPGAM | 0.97 | BRT | BRT | 0.98 | SEPGAM | SEPGAM |
| *Pseudotsuga* | 0.97 | BRT | BRT | 0.89 | SEPBRT | BRT | 0.95 | SEPGAM | BRT |
| *Quercus* | 0.78 | SEPGAM | SEPGAM | 0.73 | SEPBRT | BRT | 0.81 | SEPGAM | SEPGAM |
| *Salix* | 0.94 | BRT | BRT | 0.81 | BRT | BRT | 0.69 | SEPBRT | BRT |
| *Sapindus* | 0.94 | SEPGAM | SEPGAM | 0.999 | SEPGAM | BRT | 0.94 | SEPGAM | SEPGAM |
| *Sassafras* | 0.56 | BRT | BRT | 0.58 | SEPBRT | BRT | 0.66 | SEPGAM | BRT |
| *Sorbus* | 0.75 | BRT | BRT | 0.93 | SEPBRT | BRT | 0.86 | BRT | BRT |
| *Taxus* | 0.89 | BRT | BRT | 0.66 | BRT | BRT | 0.33 | SEPGAM | BRT |
| *Tilia* | 0.98 | SEPBRT | SEPBRT | 0.69 | BRT | SEPGAM | 0.78 | BRT | SEPBRT |
| *Tsuga* | 0.78 | BRT | BRT | 0.75 | SEPGAM | BRT | 0.6 | SEPGAM | BRT |
| *Ulmus* | 0.60 | SEPBRT | SEPBRT | 0.84 | SEPGAM | SEPGAM | 0.79 | SEPBRT | SEPBRT |

**Table S5.** Model selection results for small tree genus-specific models (n=48 genera): **a.** overall combinations and **b.** individual models for presence and number of trees. “SEP” as a prefix indicates a separate model fit to a given genus, whereas no prefix indicates a global model fit to all genera.

**a.**

| **Model types** | **Frequency** | **Proportion** |
| --- | --- | --- |
| BRT/BRT | 15 | 0.31 |
| GAM/GAM | 0 | 0 |
| SEPBRT/SEPBRT | 1 | 0.02 |
| SEPGAM/SEPGAM | 0 | 0 |
| GAM/BRT | 1 | 0.02 |
| GAM/SEPGAM | 0 | 0 |
| GAM/SEPBRT | 1 | 0.02 |
| BRT/GAM | 0 | 0 |
| BRT/SEPGAM | 12 | 0.25 |
| BRT/SEPBRT | 2 | 0.04 |
| SEPGAM/GAM | 0 | 0 |
| SEPGAM/BRT | 3 | 0.06 |
| SEPGAM/SEPBRT | 0 | 0 |
| SEPBRT/GAM | 0 | 0 |
| SEPBRT/BRT | 10 | 0.21 |
| SEPBRT/SEPGAM | 3 | 0.06 |

**b.**

| **Model type** | **Tree presence model count** | **Tree presence model proportion** | **Tree abundance model count** | **Tree abundance model proportion** |
| --- | --- | --- | --- | --- |
| BRT | 29 | 0.60 | 29 | 0.60 |
| GAM | 2 | 0.04 | 0 | 0 |
| SEPBRT | 14 | 0.29 | 4 | 0.08 |
| SEPGAM | 3 | 0.06 | 15 | 0.31 |

**Table S6.** Model selection results for medium tree genus-specific models (n=48 genera): **a.** overall combinations and **b.** individual models for presence and number of trees. “SEP” as a prefix indicates a separate model fit to a given genus, whereas no prefix indicates a global model fit to all genera.

**a.**

| **Model types** | **Frequency** | **Proportion** |
| --- | --- | --- |
| BRT/BRT | 10 | 0.21 |
| GAM/GAM | 0 | 0 |
| SEPBRT/SEPBRT | 0 | 0 |
| SEPGAM/SEPGAM | 4 | 0.08 |
| GAM/BRT | 1 | 0.02 |
| GAM/SEPGAM | 2 | 0.04 |
| GAM/SEPBRT | 0 | 0 |
| BRT/GAM | 0 | 0 |
| BRT/SEPGAM | 11 | 0.23 |
| BRT/SEPBRT | 1 | 0.02 |
| SEPGAM/GAM | 0 | 0 |
| SEPGAM/BRT | 3 | 0.06 |
| SEPGAM/SEPBRT | 0 | 0 |
| SEPBRT/GAM | 0 | 0 |
| SEPBRT/BRT | 10 | 0.21 |
| SEPBRT/SEPGAM | 0 | 0 |

**b.**

| **Model type** | **Tree presence model count** | **Tree presence model proportion** | **Tree abundance model count** | **Tree abundance model proportion** |
| --- | --- | --- | --- | --- |
| BRT | 22 | 0.46 | 29 | 0.60 |
| GAM | 3 | 0.06 | 0 | 0 |
| SEPBRT | 16 | 0.33 | 1 | 0.02 |
| SEPGAM | 7 | 0.15 | 18 | 0.38 |

**Table S7.** Model selection results for large tree genus-specific models (n=48 genera): **a.** overall combinations and **b.** individual models for presence and number of trees. “SEP” as a prefix indicates a separate model fit to a given genus, whereas no prefix indicates a global model fit to all genera.

**a.**

| **Model types** | **Frequency** | **Proportion** |
| --- | --- | --- |
| BRT/BRT | 3 | 0.063 |
| GAM/GAM | 0 | 0 |
| SEPBRT/SEPBRT | 3 | 0.063 |
| SEPGAM/SEPGAM | 5 | 0.10 |
| GAM/BRT | 3 | 0.063 |
| GAM/SEPGAM | 0 | 0 |
| GAM/SEPBRT | 0 | 0 |
| BRT/GAM | 0 | 0 |
| BRT/SEPGAM | 2 | 0.042 |
| BRT/SEPBRT | 2 | 0.042 |
| SEPGAM/GAM | 1 | 0.021 |
| SEPGAM/BRT | 13 | 0.27 |
| SEPGAM/SEPBRT | 2 | 0.042 |
| SEPBRT/GAM | 0 | 0 |
| SEPBRT/BRT | 14 | 0.29 |
| SEPBRT/SEPGAM | 0 | 0 |

**b.**

| **Model type** | **Tree presence model count** | **Tree presence model proportion** | **Tree abundance model count** | **Tree abundance model proportion** |
| --- | --- | --- | --- | --- |
| BRT | 7 | 0.15 | 33 | 0.69 |
| GAM | 3 | 0.06 | 1 | 0.02 |
| SEPBRT | 17 | 0.35 | 7 | 0.15 |
| SEPGAM | 21 | 0.44 | 7 | 0.15 |

**Table S8.** Distribution of pest-host mortality risk across severity categories (normalized for equal contribution of each pest species). **a.** Distribution across feeding guilds, including pests listed as low-impact in [7] (**A**), pest-host combinations listed as intermediate impact within [7] that were absent from [2] **(B)**, and the 6 severity categories from [2] (**C-H)**. **b.** Distribution of non-negligible impact pest species (**B-H**). **c.** Distribution of non-negligible impact pest species across host tree genera (all genera not listed are not impacted by any pest in our dataset).

| **Severity** | **A** | **B** | **C** | **D** | **E** | **F** | **G** | **H** |
| --- | --- | --- | --- | --- | --- | --- | --- | --- |
| **a. Feeding guild** |  |  |  |  |  |  |  |  |
| Borers | 71 | 15.83 | 1.89 | 2.97 | 3.33 | 4.33 | 1.50 | 2.50 |
| Defoliators | 155 | 5.54 | 3.19 | 4.00 | 1.33 | 0 | 0 | 0 |
| Sap-feeders | 192 | 6.20 | 11.95 | 4.86 | 0 | 0.30 | 0 | 0.29 |
| **b. Pest species** |  |  |  |  |  |  |  |  |
| *Adelges piceae* | 0 | 0.29 | 0 | 0.29 | 0 | 0.43 | 0 | 0 |
| *Adelges tsugae* | 0 | 0 | 0 | 0 | 0 | 0 | 0 | 1.00 |
| *Agrilus planipennis* | 0 | 0 | 0 | 0 | 0 | 0.20 | 0.30 | 0.50 |
| *Anoplophora glabripennis* | 0 | 0 | 0.13 | 0.20 | 0.33 | 0.33 | 0 | 0 |
| *Asterolecanium variolosum* | 0 | 0.77 | 0 | 0.23 | 0 | 0 | 0 | 0 |
| *Carulaspis juniperi* | 0 | 0 | 1.00 | 0 | 0 | 0 | 0 | 0 |
| *Coleophora laricella* | 0 | 0 | 0 | 0.50 | 0.50 | 0 | 0 | 0 |
| *Cryptorhynchus lapathi* | 0 | 0 | 1.00 | 0 | 0 | 0 | 0 | 0 |
| *Cyrtepistomus castaneus* | 0 | 0 | 1.00 | 0 | 0 | 0 | 0 | 0 |
| *Enarmonia formosana* | 0 | 0 | 0 | 1.00 | 0 | 0 | 0 | 0 |
| *Eulecanium cerasorum* | 0 | 0 | 1.00 | 0 | 0 | 0 | 0 | 0 |
| *Fenusa pusilla* | 0 | 0.50 | 0.33 | 0.17 | 0 | 0 | 0 | 0 |
| *Fiorinia externa* | 0 | 0.50 | 0 | 0.50 | 0 | 0 | 0 | 0 |
| *Homadaula anisocentra* | 0 | 0.50 | 0.50 | 0 | 0 | 0 | 0 | 0 |
| *Lepidosaphes ulmi* | 0 | 0.44 | 0.19 | 0.37 | 0 | 0 | 0 | 0 |
| *Lymantria dispar* | 0 | 0 | 0.05 | 0.95 | 0 | 0 | 0 | 0 |
| *Maconellicoccus hirsutus* | 0 | 0 | 1.00 | 0 | 0 | 0 | 0 | 0 |
| *Matsucoccus matsumurae* | 0 | 0 | 0 | 1.00 | 0 | 0 | 0 | 0 |
| *Neodiprion sertifer* | 0 | 0.97 | 0.03 | 0 | 0 | 0 | 0 | 0 |
| *Operophtera brumata* | 0 | 0 | 1.00 | 0 | 0 | 0 | 0 | 0 |
| *Otiorhynchus sulcatus* | 0 | 0 | 1.00 | 0 | 0 | 0 | 0 | 0 |
| *Popillia japonica* | 0 | 0.56 | 0.44 | 0 | 0 | 0 | 0 | 0 |
| *Pristiphora erichsonii* | 0 | 0 | 0 | 0.50 | 0.50 | 0 | 0 | 0 |
| **c. Host Genus** |  |  |  |  |  |  |  |  |
| *Abies* | 0 | 0.29 | 0 | 0.29 | 0 | 0.43 | 0 | 0 |
| *Acer* | 0 | 4.43 | 1.61 | 0 | 0 | 1.67 | 0 | 0 |
| *Aesculus* | 0 | 0 | 0 | 0 | 0.13 | 0 | 0 | 0 |
| *Amelanchier* | 0 | 0 | 0 | 0.50 | 0 | 0 | 0 | 0 |
| *Betula* | 0 | 0.10 | 0.13 | 0.14 | 3.00 | 0 | 0 | 0 |
| *Castanea* | 0 | 0 | 1.00 | 0 | 0 | 0 | 0 | 0 |
| *Chamaecyparis* | 0 | 0 | 1.00 | 0 | 0 | 0 | 0 | 0 |
| *Crataegus* | 0 | 0 | 1.50 | 2.50 | 0 | 0 | 0 | 0 |
| *Fraxinus* | 0 | 0 | 0 | 3.33 | 0 | 1.00 | 1.50 | 2.50 |
| *Gleditsia* | 0 | 0.06 | 0.12 | 0 | 0 | 0 | 0 | 0 |
| *Juniperus* | 0 | 0 | 0.71 | 0 | 0 | 0 | 0 | 0 |
| *Larix* | 0 | 0 | 0 | 0.24 | 0.18 | 0 | 0 | 0 |
| *Liquidambar* | 0 | 0 | 0.60 | 0 | 0 | 0 | 0 | 0 |
| *Morus* | 0 | 0 | 1.00 | 0 | 0 | 0 | 0 | 0 |
| *Ostrya* | 0 | 0 | 0 | 0.03 | 0 | 0 | 0 | 0 |
| *Pinus* | 0 | 29.00 | 1.00 | 0.17 | 0 | 0 | 0 | 0 |
| *Populus* | 0 | 0 | 0.43 | 1.00 | 0 | 0 | 0 | 0 |
| *Quercus* | 0 | 13.50 | 0 | 6.50 | 0 | 0 | 0 | 0 |
| *Salix* | 0 | 0 | 0.09 | 0.10 | 0 | 0 | 0 | 0 |
| *Sorbus* | 0 | 0 | 0 | 0.06 | 0 | 0 | 0 | 0 |
| *Taxus* | 0 | 0 | 0.14 | 0 | 0 | 0 | 0 | 0 |
| *Tilia* | 0 | 0 | 0.29 | 1.00 | 0 | 0 | 0 | 0 |
| *Tsuga* | 0 | 0.67 | 0 | 0.67 | 0 | 0 | 0 | 0.11 |
| *Ulmus* | 0 | 0.57 | 0 | 6.29 | 0 | 0 | 0 | 0 |

**Table S9.** Latin hypercube sampling parameters. 1000 samples were taken for each parameter over a uniform distribution.

| **Parameter** | **Minimum** | **Maximum** |
| --- | --- | --- |
| beta *a* | 0001 | 1.00001 |
| beta *b* | 0.01 | 2.01 |

**Table S10.** Posterior distributions of the parameters of the beta distribution (*a* and *b*), the three fitted thresholds for severity categories (*AT, BT, GT*), and of the proportional mortality within each severity category (*A-H*).

|  | **Posterior mean** | **Posterior median** | **Lower 95% Bayesian CI** | **Upper 95% Bayesian CI** |
| --- | --- | --- | --- | --- |
| *a* | 0.013 | 0.0127 | 0833 | 0.0191 |
| *b* | 0.988 | 0.938 | 0.474 | 1.793 |
| *AT* | 000211 | 0000795 | 1.16E-07 | 00125 |
| *BT* | 0427 | 0424 | 0161 | 0719 |
| *GT* | 0.969 | 0.967 | 0.952 | 0.9922 |
| *A* | 6.28E-06 | 4.36E-10 | 5.80E-33 | 7.32E-05 |
| *B* | 00267 | 00228 | 5.25E-05 | 00684 |
| *C* | 0327 | 0239 | 00389 | 0929 |
| *D* | 0.0403 | 0.0331 | 0.0107 | 0.0943 |
| *E* | 0.164 | 0.159 | 0.0245 | 0.103 |
| *F* | 0.555 | 0.527 | 0.26 | 0.93 |
| *G* | 0.964 | 0.963 | 0.951 | 0.983 |
| *H* | 0.99 | 0.991 | 0.972 | 0.9998 |

**Table S11.** Predicted tree mortality and annualized costs across land types and scenarios. These costs are only for tree removal and replacement for dead trees, and do not consider non-treatment costs such as property value losses as examined in [7] or any ecosystem services losses. Community trees are defined as all urban trees apart from street trees (e.g. parks, industrial areas etc.), while residential trees are the responsibility of homeowners and are on their properties. Mean mortality for community trees in the reasonable scenario was 1.2% (83.4M trees), with an estimated annualized management cost of 1.5B USD (32.8B from 2020 to 2050), and mean mortality for residential trees was 1.1% (15.5M trees), corresponding to an annualized estimated management cost of 356M USD (8.0B from 2020 to 2050).

| **Land Type** |  | **Annualized Cost (millions 2019 USD)** | | **Tree Mortality (millions)** | | | **Total tree abundance (millions)** | **Percent mortality** | |
| --- | --- | --- | --- | --- | --- | --- | --- | --- | --- |
|  | **Mortality Debt** | **lower 95% CI** | **upper 95% CI** | | **lower 95% CI** | **upper 95% CI** |  | **upper 95% CI** | **lower 95% CI** |
| Community |  |  |  | |  |  |  |  |  |
|  | Reasonable | 1300 | 1707 | | 74.3 | 97.0 | 7000 | 1.1% | 1.4% |
|  | All 10 | 1096 | 1276 | | 62.4 | 72.5 | 7000 | 0.9% | 1.0% |
|  | All 50 | 2581 | 3202 | | 147 | 181 | 7000 | 2.1% | 2.6% |
|  | All 100 | 5618 | 6568 | | 320 | 373 | 7000 | 4.6% | 5.3% |
|  | Vary Borers | 1195 | 6279 | | 68.0 | 356 | 7000 | 1.0% | 5.1% |
|  | Vary Defoliators | 1159 | 1508 | | 64.7 | 94.8 | 7000 | 0.9% | 1.4% |
|  | Vary Sap-Feeders | 1136 | 1667 | | 66.0 | 85.4 | 7000 | 0.9% | 1.2% |
| Residential | Reasonable | 315 | 416 | | 13.7 | 15.4 | 1400 | 1.0% | 1.1% |
|  | All 10 | 264 | 308 | | 11.5 | 13.4 | 1400 | 0.8% | 1.0% |
|  | All 50 | 623 | 777 | | 27.1 | 33.6 | 1400 | 1.9% | 2.4% |
|  | All 100 | 1357 | 1593 | | 59.0 | 69.0 | 1400 | 4.2% | 4.9% |
|  | Vary Borers | 288 | 1520 | | 12.5 | 65.8 | 1400 | 0.9% | 4.7% |
|  | Vary Defoliators | 279 | 366 | | 12.1 | 15.8 | 1400 | 0.9% | 1.1% |
|  | Vary Sap-Feeders | 274 | 406 | | 11.9 | 17.6 | 1400 | 0.9% | 1.3% |

**Table S12.** List of risk factors for new high impact invaders

| **Risk Characteristic** | **Value** |
| --- | --- |
| Feeding Guild | Wood Borer |
| Preferred Hosts | *Acer, Quercus* |
| Mortality | Category H (98.98% ca. EAB) |
| Mortality Debt | 10 year (ca. EAB) |
| Continent of origin | Asia |

**Table S13.** We used the list of risk factors in **Table S11** to simulate dispersal from three of the top 4 major US ports (by total trade volume, <http://aapa-ports.org>) to forecast potential *Acer* (maple) and *Quercus* (oak) mortality and associated costs. We found that damages were highest to maple and oak trees when pests were introduced via the port of South Louisiana (SL), and lowest when introduced to the Port of New York and New Jersey (NY). The Port of Long Beach (LA) had intermediate results.

|  | **Port** | **NY** | **LA** | **SL** |
| --- | --- | --- | --- | --- |
| **Genus** |  |  |  |  |
| Maple | **Exposure** | 3.12E+06 | 2.56E+06 | 4.27E+06 |
|  | **Mortality** | 3.09E+06 | 2.54E+06 | 4.23E+06 |
|  | **Cost ($US)** | 2.37E+09 | 1.91E+09 | 3.19E+09 |
| Oak | **Exposure** | 8.12E+05 | 8.12E+05 | 1.85E+06 |
|  | **Mortality** | 8.05+05 | 8.13E+05 | 1.83E+06 |
|  | **Cost ($US)** | 7.31E+08 | 7.39E+08 | 1.66E+09 |

**Table S14.** Visual representation of the sources of uncertainty and their relative quantification across impact types, where redder hues indicate greater uncertainty. Sources of uncertainty include 1) future climatic variability, 2) spread model uncertainty, 3) host distributional model uncertainty, 4) asymptotic mortality misestimation, 5) mortality debt model misspecification, 6) variability in management behavior. Our models also do not account for the preventative cutting after the last street tree inventory date, which affected many cities across IN, IL, MI, and WI. Preventative cutting would have led to the payment of tree removal costs prior to our estimation window. This is particularly likely to have inflated the 2020-2050 costs to communities with large street tree budgets in regions where EAB was predicted to invade in the years 2010-2020 (therefore initiating mortality 2020-2030).

| **Tree Type** | **Dimension of Impact** | | | |
| --- | --- | --- | --- | --- |
|  | Exposure^1,2,3^ | Mortality (Asymptotic)^1,2,3,4^ | Mortality (2020-2050)^1,2,3,4,5^ | Cost (2020-2050)^1,2,3,4,5,6^ |
| Residential |  |  |  |  |
| Community |  |  |  |  |
| Street |  |  |  |  |

Dataset S1 (separate file). The strength of model fit across all 16 possible model combinations for each host tree genus. “SEP” as a prefix indicates a separate model fit to a given genus, whereas the lack of such a prefix indicates a global model fit to all genera. Some genera were very rare within our inventoried communities at a given size class, and therefore their genus-specific models did not have sufficient data to be fit. R^2^ is reported as NA in these cases.
